# ZAKα is a sensor of mRNA stasis at the ribosomal exit channel

**DOI:** 10.1101/2025.11.22.689755

**Authors:** Anna Constance Vind, José Francisco Martínez, Zhenzhen Wu, Andrii Bugai, Kelly Mordente, Qiuyan Chen, Mads Rasmussen, Dandan He, Jesper Q. Svejstrup, Torben Heick Jensen, Melanie Blasius, Simon Bekker-Jensen

**Author notes:** Correspondence and requests for materials should be addressed to S.B.-J. Equal contribution (Anna Constance Vind, José Francisco Martínez, Zhenzhen Wu).

## Abstract

Despite a growing interest in the ribotoxic stress response (RSR), it remains unknown how the upstream p38 and JNK-activating MAP3 kinase ZAKα senses translational impairment. Combining Alphafold3 prediction and RNA crosslinking and immunoprecipitation (CLIP), we uncover that ZAKα dynamically monitors the mRNA exit channel of elongating ribosomes for mRNA stasis. This is accomplished by ZAKα via its direct interactions with the ribosomal proteins RACK1 and RPS27 as well as with the 18S rRNA helix-26. In this conformation, four mRNA-binding peptides in ZAKα span across the path of ribosome-exiting mRNA. Progressive elongation effectively threads ZAKα off the ribosome, while mRNA stasis stabilizes the interaction allowing for kinase activation. Prolonged binding of ZAKα to slow-elongating, stalled and collided ribosomes is associated with sequestration of the inhibitory SAM domain on RACK1, allowing for transient ZAKα dimerization, activation loop trans-autophosphorylation and RSR activation. We propose that compromised ribosome processivity constitutes a common ribotoxic stress signal and that ZAKα is a ribosome collision-agnostic sensor of such perturbations.

## Introduction

The ribotoxic stress response (RSR) denotes a cellular stress response pathway in which the MAP3 kinase ZAKα senses translational aberrations and signals through p38 and JNK kinases (*1*). RSR signaling thus holds a potential to impact directly on stress response outcomes such as cell cycle arrest, programmed cell death and inflammation. RSR-activating insults can be sustained by a plethora of environmental and endogenous stressors (e.g. UV-irradiation (*2*), reactive oxygen species (*3*)), plant and microbial toxins (e.g. ricin, anisomycin (*4*)) and can even be inflicted on purpose by cellular enzymes (e.g. SLFN11 (*5*), RNase L (*6, 7*)). Recently, the RSR pathway has been shown to be critically important for a range of biologically important phenomena, including, but not limited to, UV radiation-induced cell death (*8–10*), dissemination of *Legionella* bacteria from infected cells (*11*), sensitivity of cancer cells to chemotherapy (*5*), endoreplication during tumorigenesis (*12*), UV-induced skin inflammation (*8*) and metabolic regulation in obesity and aging (*3*). The RSR is just one of several highly studied translation surveillance pathways which also include the GCN2-activated integrated stress response (ISR), ZNF598-activated ribosome-associated quality control (RQC), RNF10-activated 40S subunit degradation and an emerging GCN1-dependent ribosome ubiquitination cascade (*13–15*).

Despite the recent surge in RSR research, it remains unknown how the proximal component, the ZAKα kinase, senses translational impairment to mediate these diverse responses. ZAKα has been posited to specifically recognize collided ribosomes (*16*), which is a common hallmark of translational stress (*17, 18*). In this scenario, ZAKα would need to associate with two ribosomes simultaneously or recognize the collision interface to gain specificity in ribotoxic stress sensing.

Several well-established examples of signaling from collided ribosomes from eukaryotes (*19–21*) and bacteria (*22–24*) lend inspirational support to this notion. ZAKα has also been posited to recognize single stalled ribosomes (*3, 25*), supported by an apparent lack of correlation between the cellular amounts of collided ribosomes and RSR signaling output. Unlike other translation surveillance sensors (GCN2, ZNF598, RNF10), ZAKα appears to be dynamically associated with elongating ribosomes (*26*), surveilling the translation process in a continuous scanning fashion.

Given the propensity for rapid and potent RSR activation and the strong imbalance between the number of ZAKα molecules (app. 15,000 per cell) and ribosomes (1-10 mio. per cell), this interaction must be highly transient and have a high off rate during unperturbed translation. In addition to an N-terminal kinase domain, ZAKα harbors three structured domains; a leucine zipper (LZ), a sterile alpha-motif (SAM) and a YEATS-like domain (YLD) with unknown functions. Ribosome binding has been shown to depend on the C-terminal domain (CTD) which together with the Sensor (S) domain is redundantly required for ZAKα activation by all known ribotoxic stress signals (*26*). Both the CTD and S domains reside in the unstructured C-terminus of ZAKα that likely mediates transient interactions with the ribosome.

The current notion in the field is that ZAKα can detect both stalled and collided ribosomes with a preference for the latter. This begs several questions related to the identity of the elusive ribotoxic stress signal(s) that remain unanswered and precludes a proper understanding of the signal-sensor relationship between the ribosome and ZAKα. Here we provide an explanation for how ZAKα can respond to a plethora of translation-perturbing insults by recognizing a single common feature of elongation-impaired ribosomes. ZAKα binds to the ribosome in the vicinity of the mRNA exit channel and further attaches to the exiting mRNA via four short peptides. As the multiple binding sites for ZAKα on mRNA and the ribosome naturally separate from each other during elongation, ribosomal processivity inversely correlates with binding time. In this scenario, ZAKα is a collision-agnostic sensor of mRNA stasis in the exit channel. We propose that this new insight rationalizes diverging opinions in the translation field regarding how the RSR is activated by highly diverse insults and ribosomal conformations.

## Results

### RACK1 is critical for ZAKα ribosome interaction and activation

Likely due to the highly dynamic nature of the interaction between ZAKα and the ribosome, it has so far proven challenging to glean structural insight into this complex from cryogenic electron microscopy (cryo-EM) or related techniques. To explore alternatives, we used Alphafold3 (AF3) (*27*) modelling to predict potential interactions between full-length human ZAKα and all proteins associated with the term “translation” in Reactome (*28*) (reactome.org - R-HSA-72766). This protein set included all of the known ribosomal proteins and known translation factors. AF3 predicted with high confidence (ipTM > 0.8) an interaction with the ribosomal protein RACK1 and with somewhat lower confidence (0.8 > ipTM > 0.6) RPS27 and its paralog RPS27L (Fig. 1a; Fig. S1a). We then extracted the maximal ZAKα interaction probability (from 0 to 1) from AF3 for each amino acid from all ribosomal proteins and painted it onto a structure of the human ribosome (PDB 4UG0 (*29*)). This analysis highlighted exclusively RACK1 (two surface-exposed patches) and RPS27 (one surface-exposed patch) as potential protein-protein interaction sites for ZAKα on the ribosome and in relatively close proximity to each other (Fig. 1b). By revisiting a historical genetic screen reporting on ZAKα activation (*30*), we noticed that the semi-essential *RACK1* consistently scored as a strong positive regulator while *RPS27* provided a weaker and less significant hit (Fig. 1c). We could validate the requirement of RACK1 for anisomycin-induced ZAKα activation in both HAP1 and Hela cells deleted for *RACK1,* and this effect was fully rescued by re-introduction of ectopic RACK1 (Fig. 1d; Fig. S1b,c). Furthermore, even though RACK1-deficient ribosomes from these cells could both translate and collide (Fig. S1c,d), they did not co-purify ZAKα, suggesting that RACK1 is indeed critical for ribosome binding of the RSR-activating kinase (Fig. 1d; Fig. S1b). To further understand the structural basis of this interaction, we examined the AF3-provided dimeric complexes consisting of ZAKα / RACK1 and ZAKα / RPS27. The resulting PAE matrices and inspection of the proposed structures suggested that the interactions are based on three short linear interaction motifs (SLIMs) with a length of 5-6 amino acids. Two of these reside in the unstructured C-terminal 200 amino acids of ZAKα, while the second RACK1-binding motif is located in close proximity to the SAM domain (Fig. 1e-i). A virtually identical mode of binding to the same two RACK1 pockets via two SLIMs (“CR1” and “CR2”) was recently described for the translation-regulating factors LARP4 and LARP4B (*31*). The two RACK1-binding SLIMs in ZAKα are almost identical to these sequences (Fig. S1e), suggesting a conserved mode of ribosome interaction. Deletion of either of the two SLIMs predicted to bind RACK1 (ZAKα Δ417-422 and Δ611-617) rendered ZAKα refractory to activation by anisomycin (Fig. 1j), underscoring the critical nature of both interactions. While the ZAKα Δ417-422 mutant retained the ability to co-purify with ribosomes, deletion of the other RACK1-binding SLIM (ZAKα Δ611-617) completely abrogated this interaction (Fig. 1k), consistent with the requirement of the RACK1 protein for the same (Fig. 1d; Fig. S1b). Deletion of the predicted RPS27-binding SLIM alone (ZAKα Δ768-772) did not impair ZAKα activation by anisomycin (Fig. 1l), but we did note a reduced binding of the last 200 amino acids of ZAKα to the ribosome when this motif was deleted (Fig. S1f). Our data thus shows that RACK1 is a critical ribosomal hub for both ribosome binding and activation of ZAKα.

**Figure 1.**
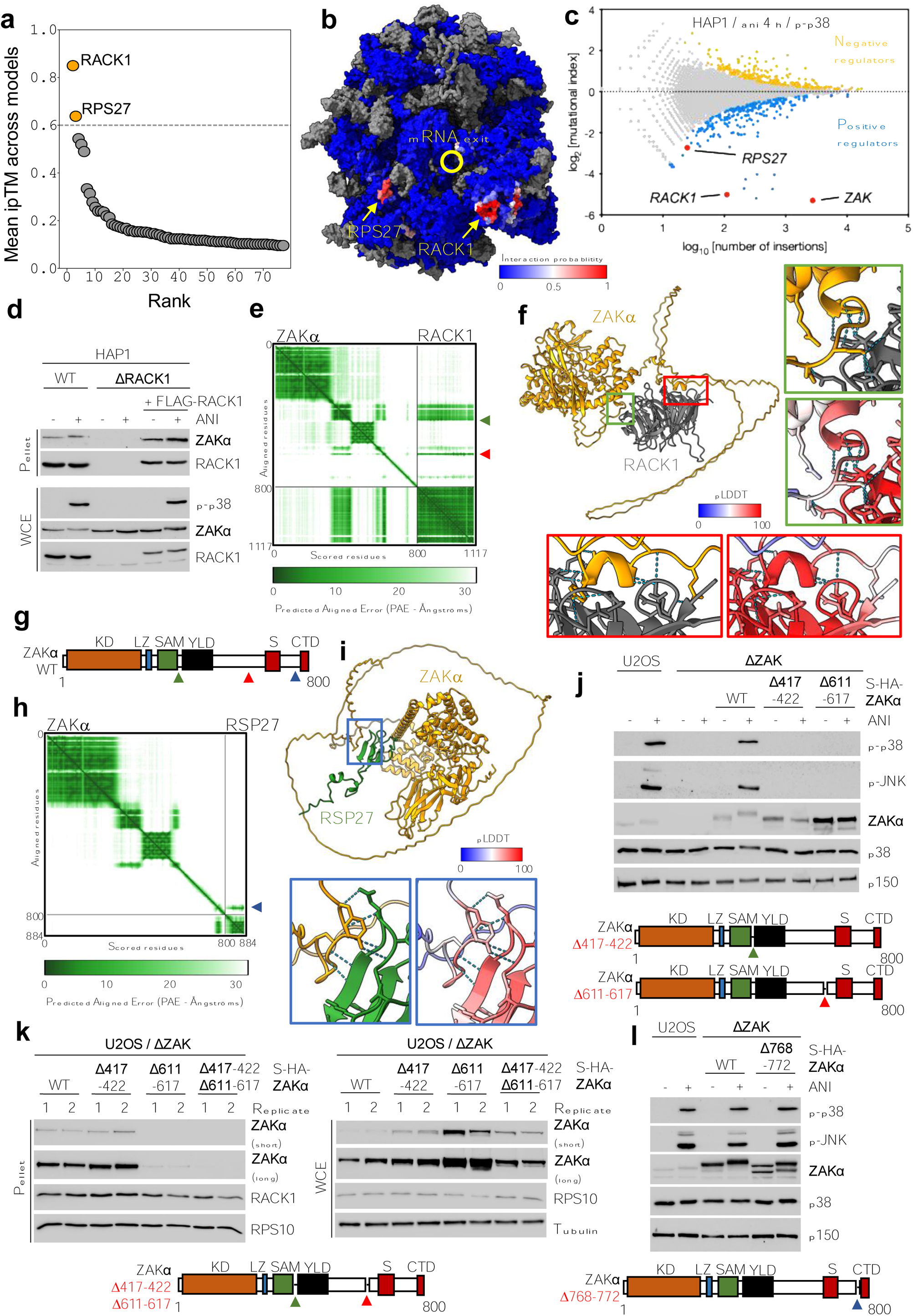
Alphafold modelling identifies ribosomal protein components binding to ZAKα. **a.** Alphafold3 (AF3) prediction scores for ZAKα binding to individual ribosomal proteins, ranked according to ipTM score. Dashed line indicates our cutoff score of ipTM = 0.6. **b.** Predictions from (a) were used to calculate per-residue interaction probabilities. Values (0/blue to 1/red) were used to paint a structure of the human ribosome (PDB 4UG0). **c.** Gene-trap-based genetic screen in haploid human cells for regulators of p38 kinase activation in response to anisomycin (ani). Data re-analysed from (*30*) with predicted binding partners *RACK1* and *RPS27* highlighted. **d.** HAP1 WT, ΔRACK1 and ΔRACK1 cells stably rescued with FLAG-tagged RACK1 were treated with ani (1 μM - 1 h) and lysates were ultracentrifugated through a sucrose cushion. Whole cell extract (WCE) and pelleted material (pellet) enriched for ribosomes were analyzed by immunoblotting with the indicated antibodies. **e.** Predicted aligned error (PAE) matrix plot of an AF3-generated prediction of a ZAKα-RACK1 interaction structure. **f.** AF3-generated structure from (e). Blow-ups of the two interaction sites (also indicated by green and red arrows in (e)) with hydrogen bonds between chains indicated and colored according to the predicted local distance difference test (pLDDT) score. **g.** Domain structure of ZAKα protein with RACK1-binding (green and red arrows) and RPS27-binding (blue arrow) regions indicated. KD, kinase domain; LZ, leucine zipper; SAM, sterile alpha-motif; YLD, Yeats-like domain; S, sensor domain; CTD, C-terminal domain. **h.** PAE matrix plot of an AF3-generated prediction of a ZAKα-RPS27 interaction structure. **i.** AF3-generated structure from (h). Blow-up of the interaction site (also indicated by blue arrow in (h)) displayed as in (f). **j.** U2OS / ΔZAK cells stably rescued with WT and mutated forms of strep-HA-tagged ZAKα were treated as in (d) and analyzed by immunoblotting with the indicated antibodies. **k.** U2OS / ΔZAK cells stably rescued with WT and mutated forms of strep-HA-tagged ZAKα were analyzed for ribosome interaction as in (d) with duplicate samples. **l.** As in (j), except that cells were rescued with a mutant of ZAKα predicted to be deficient for RPS27 binding.

### iCLIP highlights an rRNA component of the ZAKα-ribosome interaction

We found it plausible that ZAKα engages both protein and rRNA components when transiently contacting the ribosome. While AF3 modelling successfully predicted the former, this tool is of little use in predicting the latter. Instead, we employed individual-nucleotide resolution crosslinking and immunoprecipitation (iCLIP) technology (*32*) to map rRNA bases in close proximity to ZAKα (Fig. 2a,b; Fig. S1g,h). We compared rRNA crosslink of strep-HA-tagged WT ZAKα in the absence or presence of low (L) or high (H) anisomycin for 15 minutes, conditions that are believed to distinguish collision (L) from stalling / ”freezing” (H) of individual ribosomes (*16, 33*).

**Figure 2.**
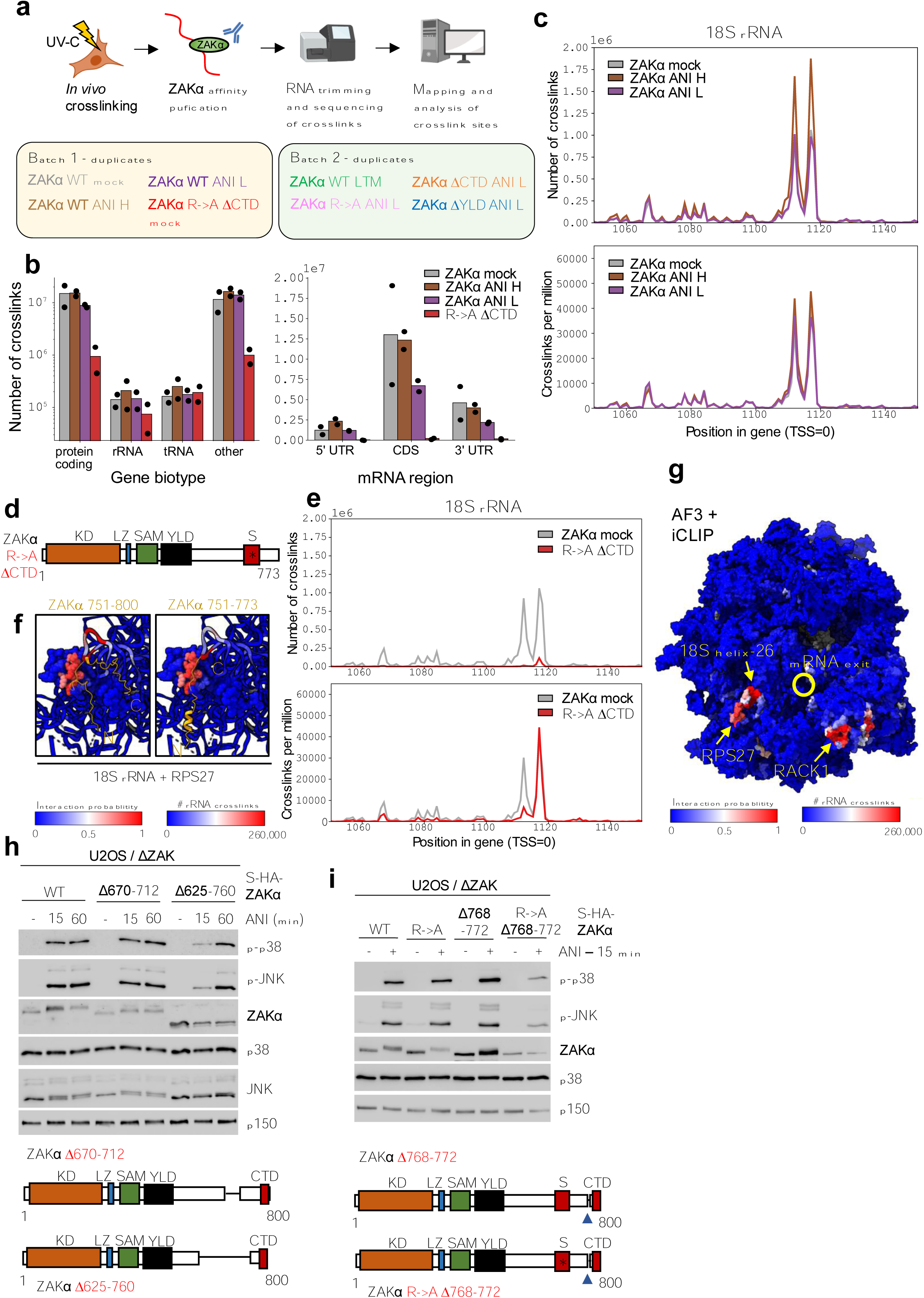
iCLIP reveals an RPS27-proximal rRNA binding site for ZAKα. **a.** Schematic of individual-nucleotide resolution crosslinking and immunoprecipitation (iCLIP) protocol and overview of biological duplicate samples sequenced in two batches. **b.** Left: Total number of ZAKα crosslinks from duplicate samples of sequencing batch 1 according to RNA category (gene biotype). Right: As in (left), except according to mRNA elements. **c.** Total number of crosslinks (top) and normalized rRNA crosslink counts (bottom) for individual nucleotides in a region of 18S rRNA. Mock treatment, low anisomycin (ani L – 0.19 μM) and high anisomycin (ani H - 76 μM) (15 min) conditions have been overlaid. The region represents the most prominent crosslink peak from the analysis in Fig. S2a. Values represent the mean of duplicate experiments. **d.** Schematic of a ribosome binding and activation-deficient mutant of ZAKα (R->A ΔCTD). **e.** As in (c), except that total and normalized crosslinks (normalized by library size) for ZAKα WT vs R->A ΔCTD was analyzed. Excerpt from Fig. S2c. **f.** Structural alignment of an experimental structure of the human ribosome (PDB 4UG0) with AF3-predicted complexes of RPS27 and the last 50 amino acids of ZAKα with (left) or without (right) the CTD included. Notice that CTD-deleted ZAKα can still contact part of 18S helix-26 when bound to RPS27. RPS27 in 4UG0 (other ribosomal proteins are hidden) is colored according to interaction probability (protein - AF3 prediction) and total number of WT ZAKα crosslinks (rRNA). **g.** 4UG0 painted by per-residue ZAKα interaction probabilities from Fig. 1b was further painted with total number of ZAKα crosslinks per rRNA residue. Combining prediction of protein interaction (AF3) and experimental determination of rRNA crosslink sites (iCLIP) highlights two ribosomal binding patches for ZAKα in close proximity to the mRNA exit channel. Structural ribosomal elements involved in the interaction are highlighted in yellow. **h.** U2OS / ΔZAK cells stably rescued with strep-HA-tagged ZAKα mutants with shortened linker (Δ670-712 and Δ625-760) were treated with anisomycin (Ani – 1 μM) for the indicated times **i.** As in (h), except that mutants deficient for RPS27 binding and/or S domain functionality were treated with ani (1 μM) for 15 min. TSS, transcription start site.

Remarkably, the number of sequenced crosslinks were orders of magnitude higher for 18S rRNA than for any of the three other rRNAs (Fig. S2a), supporting that ZAKα binds the 40S subunit of the ribosome. A very prominent double-peak centered at U1114 and U1120 (Fig. 2c) caught our attention as this corresponds to the 18S helix-26 that is placed immediately adjacent to the AF3-proposed ZAKα-RPS27 binding site in the ribosome structure. To control for the validity of these crosslinks, we also analyzed our previously described double S and CTD mutant of ZAKα (ZAKα R->A ΔCTD (Fig. 2d; Fig. S2b)) that is refractory to both ribosome binding and activation (*26, 34*). While this mutant returned as many crosslinks to 28S, 5S and 5.8S rRNA as WT ZAKα, the amount of crosslinking to 18S rRNA was strongly reduced (Fig. 2e – top; Fig. S2c). After normalization (crosslinks per million crosslinks), the crosslinking efficiency of the ZAKα mutant, especially to U1120 however still remained considerable (Fig. 2e – bottom), demonstrating that its rRNA binding capacity *per se* was not impaired. Structural alignment of the ribosome with an AF3-generated structure of RPS27 complexed with the last 50 amino acids of ZAKα (+/- CTD) was consistent with this mutant still being able to contact RPS27 and crosslink to the adjacent nucleotides of 18S helix-26 (Fig. 2f). Combining AF3 modelling and iCLIP analysis by simultaneously color-coding the ribosomal surface for AF3-based interaction probabilities (Fig. 1b) and iCLIP-based crosslink counts highlighted how ZAKα engages two sites on either side of the ribosomal mRNA exit channel (Fig. 2g). Inspection of this combined model of ribosome-ZAKα interaction from all angles did not point to any alternative interpretation of our data (Fig. S2d). Importantly, the overall iCLIP profile was largely unchanged upon addition of either concentration of anisomycin (Fig. 2c; Fig. S2a), suggesting that interaction with this ribosomal surface is equally relevant for the scanning and sensing modes of ZAKα on the ribosome. We investigated the importance of this dual interaction mode by generating and testing several deletion mutants of ZAKα: Firstly, we shortened the linker between the second RACK1-binding SLIM and the RPS27-binding SLIM to an extent where ZAKα cannot simultaneously occupy both binding sites (ZAKα Δ625-712) and secondly, we combined the deletion of the RPS27-binding SLIM with amino acid substitutions at multiple positions of the S domain (ZAKα R->A Δ768-772). These mutants were quite capable of activation when assayed 1 hour after anisomycin treatment (Fig. 2h; S3a). However, they were also clear hypomorphs, being strongly compromised for activation when assayed at a shorter time (15 minutes) after exposure to anisomycin (Fig. 2h,i).

### ZAKα S and CTD domains bind mRNA to achieve sufficient affinity for ribosome interaction

Our iCLIP experiments also returned high amounts of ZAKα crosslinks on mRNA (Fig. 2b; Fig. S1h), and metagene profiles of total as well as normalized counts demonstrated that these were enriched within the 5’UTR and the coding sequence (i.e. the mRNA bases that are in direct contact with the ribosome during initiation and elongation) (Fig. 3a,b). In stark contrast to WT ZAKα, the ribosome binding-deficient ZAKα R/K->A ΔCTD mutant barely returned any mRNA crosslinks (Fig. 3a), suggesting that binding of ZAKα to mRNA correlates directly with ribosome interaction. We therefore sought to unravel the relative contributions of the ZAKα S and CTD domains to mRNA and ribosome binding. While both single mutants were refractory for binding to ribosomes during unperturbed translation, they both exhibited relatively strong ribosome interaction in anisomycin-treated cells and were able to support RSR signaling (Fig. 3c). This was in contrast to the double mutant which was refractory to both activities (Fig. 3c). A close inspection of the few mRNA crosslinks of the double mutant after normalization returned only a seeming pattern of noise that did not resemble the normalized metagene profiles of WT ZAKα across the whole mRNA or around the start and stop codons (Fig. S3b). Conversely, each of the single ZAKα mutants behaved similar to WT indicative of direct mRNA binding (Fig. S3c). These results, thus, explain the curious redundancy of the S and CTD domains (*8, 26*) in that they both contact mRNA to mediate ribosome binding and ribotoxic stress sensing.

**Figure 3.**
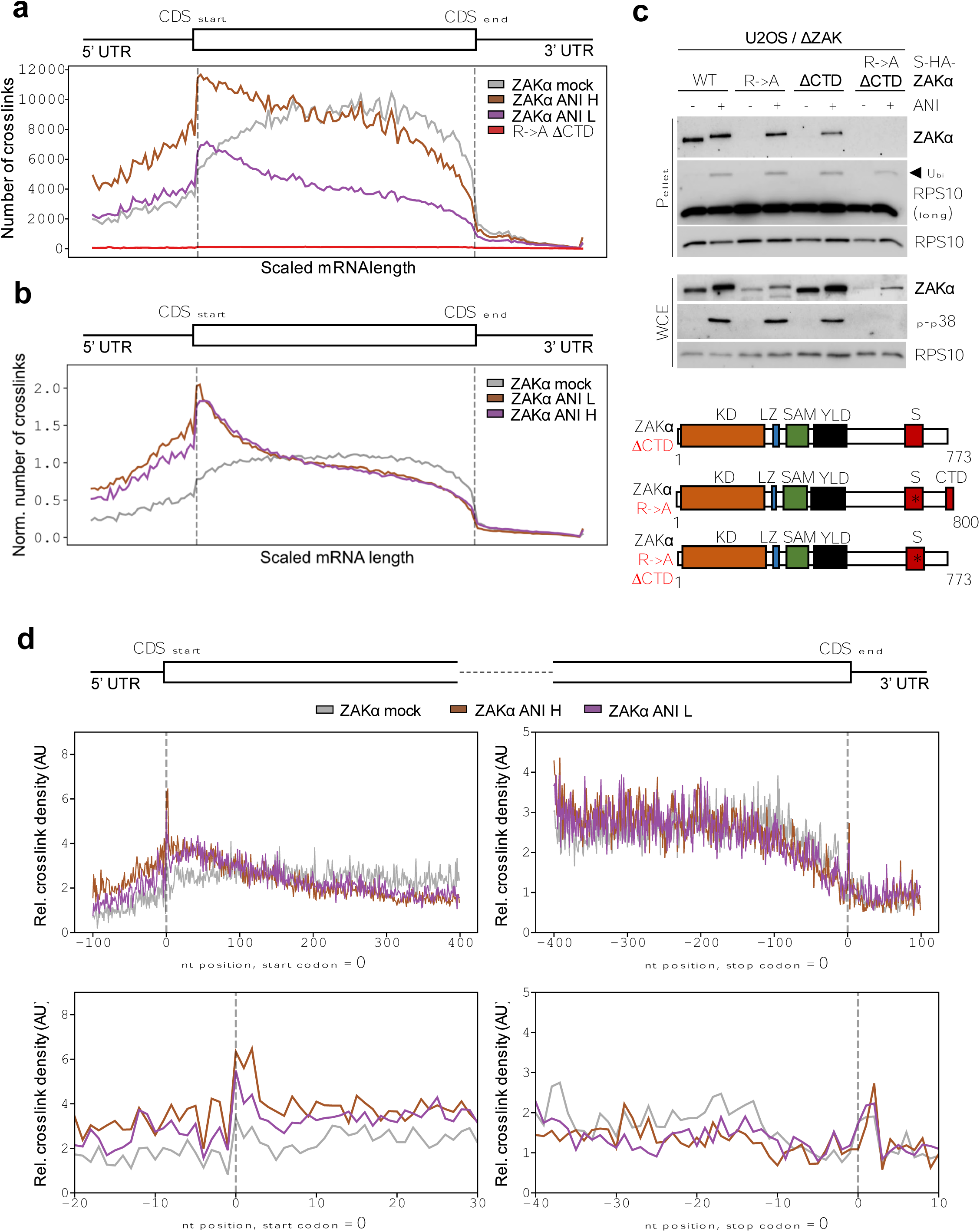
S and CTD are mRNA-binding domains critical for ZAKα-ribosome interaction. **a.** Metagene profiles of total number of crosslinks for ZAKα WT and R->A ΔCTD along scaled length of spliced mRNAs determined by iCLIP. Values represent means from biologial duplicates. **b.** As in (a), except that normalized crosslink numbers were plotted. **c.** U2OS / ΔZAK cells stably rescued with strep-HA-tagged ZAKα WT and mutants (bottom) were treated with ani (1 μM - 1 h) and lysates were ultracentrifuged through a sucrose cushion. Whole cell extract (WCE) and pelleted material (pellet) enriched for ribosomes were analyzed by immunoblotting with the indicated antibodies. **d.** Analysis of normalized mRNA crosslinks for ZAKα treated with the indicated conditions of anisomycin (ani L – 0.19 μM; ani H - 76 μM) around the start (left) and stop codons (right) shown at low (top) and high (bottom) resolution. Notice the anisomycin-induced increase of crosslinks upstream of the start codon and the apparent lack of crosslinks immediately upstream of the stop codon. nt, nucleotide number.

The normalized metagene profile of WT ZAKα across the mRNA highlighted that the bulk of binding occurred in the middle of the coding sequence (CDS) and in the 5’UTR but not in the 3’UTR (Fig. 3b). This profile changed dramatically with anisomycin treatment with a bias towards the 5’UTR and the upstream parts of the CDS and with lower occupancy around the stop codon. These profiles were however identical between low (“collision stress”) and high anisomycin (ribosome “freezing” conditions) treated samples (Fig. 3b), indicating that ZAKα does not bind mRNA or ribosomes differentially between these two conditions. Our map of ribosomal binding surfaces indicated that ZAKα surveils the mRNA exit channel of the ribosome (Fig. 2g). A closer inspection of mRNA crosslink sites around the stop codon supported this hypothesis, highlighting a sharp decrease in crosslinks in all conditions at around 12 nucleotides upstream from the stop codon (Fig. 3d - right). This gap is reminiscent of a ribosome-protected mRNA fragment originating from a ribosome occupying the last codon of the open reading frame (ORF) (*35*). A similar inspection of the start codon-proximal region revealed the presence of crosslinks in the last part of the 5’ UTR, preferentially from anisomycin-treated samples and likely represented binding to ribosomes that have been arrested at or closely after the start codon (Fig. 3d - left).

### Impaired displacement of mRNA from the ribosomal exit channel is a signal for ZAKα activation

Treatment of cells with ribosome inhibitors is associated with stalling of individual ribosomes. On mRNAs with several ribosomes engaged in translation, individual stalled ribosomes can collide with elongating ones. An often used and simple experimental way of distinguishing between ‘stalled + collided’ and ‘stalled only’ ribosomes is to treat cells with saturating amounts of ribosome inhibitors, e.g. anisomycin, which in principle should freeze all ribosomes to elicit a ‘stalling only’ response. We treated U2OS cells deleted (U2OS / ΔZAK) or not (WT) for the *ZAK* gene with increasing concentrations of anisomycin. At saturating conditions (100 μg/ml), 15 minutes of anisomycin treatment was associated with ZAKα-dependent p38 and JNK activation (Fig. 4a). Albeit this activation was somewhat less than after lower doses of anisomycin it was clearly detectable. When we extended the treatment period to 1 hour, no difference in RSR signaling could be observed across the different anisomycin concentrations (Fig. S4a). We conclude that it is not possible to abolish ZAKα activation by anisomycin over-titration and elimination of ribosome collision. We previously reported (*26*) that the E-site ribosome inhibitor lactimidomycin (LTM), which stalls exclusively initiating 80S ribosomes, is also a potent ZAKα activator (Fig. S4b). To investigate whether this activation is due to sensing of single stalled ribosomes or is dependent on collision with 40S subunits scanning the 5’ UTR, we created a cell line where the endogenous ORF of the EIF3A component of the EIF3 complex is fused to the FKBP12-F36V inducible degradation tag. Treatment of cells with the proteolysis-targeting chimera (PROTAC) compound dTAG^V^-1 led to complete degradation of EIF3A within 1 hour and an associated complete loss of new protein synthesis as measured by puromycin incorporation (Fig. 4b). We next co-treated these cells with LTM and dTAG^V^-1 for 1 hour, creating a scenario in which 40S scanning activity is absent while 80S ribosomes are still stalled at start codons. Even here strong ZAKα activation ensued (Fig. 4c), highlighting that both 80S-80S and 40S-80S collisions are dispensable for RSR signaling. iCLIP of WT ZAKα from cells treated with a low dose of LTM (1 μM) for 15 minutes (Fig. 2a; Fig. S1h) returned a large number of mRNA crosslinks upstream of the start codon (Fig. S4c). Inspection of the normalized metagene profiles for LTM-treated and unperturbed cells around the start codon revealed that this peak had its zenith at app. 9 nucleotides upstream of the start codon and did not extend into the CDS (Fig. 4d). These results, along with our AF3-modelling and other iCLIP data, allowed us to formulate a model for ribosome-binding by ZAKα. Here, ZAKα associates transiently with RACK1, RPS27, 18S helix-26 and mRNA protruding from the mRNA exit channel. On elongating ribosomes, movement of the mRNA component of this composite binding reaction will destabilize the interaction, effectively ejecting ZAKα from the ribosome shortly upon arrival (Fig. 4e). We further propose that ZAKα from this position can sense compromised processivity of stalled 80S ribosomes only when they are loaded onto mRNA. We previously reported (*25*), and here corroborated, that mTOR inhibition leads to weak ZAKα activation (Fig. 4f – compare lanes 1-2 with lanes 7-8). Apart from downregulation of cap-dependent initiation, mTOR inhibition is also associated with activation of the kinase EEF2K which negatively regulates ribosomal elongation speed (*36*). Indeed, the EEF2K-activating compound nelfinavir (*37*) was recently shown to cause EEF2K-dependent ISR activation, a phenomenon coined “elongation stress” (*38*). Addition of nelfinavir to cells also caused ZAKα-dependent RSR activation (Fig. 4f). A reduction in ribosomal elongation speed by activation of EEF2K via either an mTOR-inactivating compound or nelfinavir is thus sufficient to activate ZAKα, supporting the notion that this sensor responds to mRNA stasis and/or slow movement of mRNA through the ribosomal exit channel. We then considered whether our proposed binding and sensing mode would be consistent with simultaneous recognition of both entities of a collided ribosome. Interrogating a structure of a disome (PDB 7QVP (*20*)), it appeared unlikely that ZAKα can simultaneously connect to RACK1 and RPS27-18S helix-26 of the leading (stalled) ribosome while also gaining access to the very short stretch of mRNA bases that are partially buried at the collision interface (Fig. S4d). It is however entirely possible for ZAKα to employ the same binding and sensing mode on the colliding ribosome in the disome structure as the one we propose for single stalled ribosomes (Fig. 4e). Based on the above, we therefore suggest that ZAKα is a sensor of mRNA stasis in the exit channel and that this is independent of whether the stress signals arise due to the presence of slow-moving/paused, stalled or collided ribosomes (Fig. 4e).

**Figure 4.**
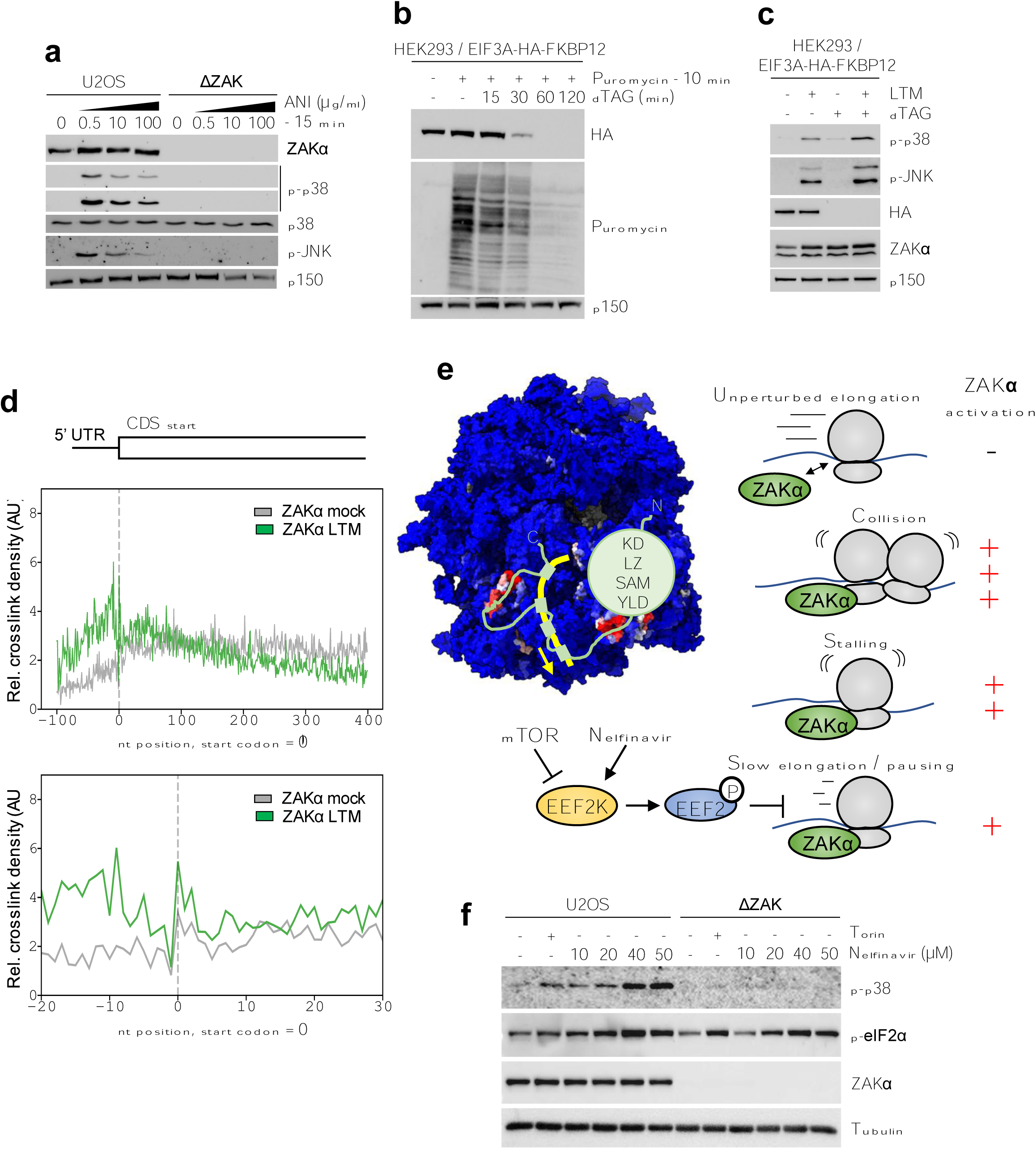
ZAKα binds to mRNA exiting the ribosome and is a sensor of impaired elongation. **a.** U2OS and ΔZAK cells were treated with increasing concentrations of anisomycin (ani – 1, 10 and 100 μg/ml) for 15 min. Lysates were analyzed by immunoblotting with the indicated antibodies. **b.** HEK293 cells with stable integration of HA-tagged FKBP12-F36V in frame with the endogenous *EIF3A* gene were induced to degrade EIF3A by treatment with dTAG^V^-1 (500 nM) for the indicated times. Translation output was determined by puromycin incorporation assay. **c.** Cells from (b) were treated with lactimidomycin (LTM, 1 μM – 1 h) and/or dTAG^V^-1 (500 nM – 1 h). Lysates were analyzed as in (a). **d.** Analysis of normalized mRNA crosslinks for ZAKα from cells treated with LTM (1 μM – 15 min) around the start codon shown at low (top) and high (bottom) resolution. nt, nucleotide number. **e.** Left: Model of ZAKα binding to translating ribosomes. The flexible C-terminus of ZAKα anchors on two ribosomal surface patches and four peptide motifs contact the mRNA that protrudes from the exit channel. The green sphere represents folded protein domains in the N-terminal part of ZAKα. Productive elongation combined with obligate mRNA interaction will result in rapid separation of binding sites and displacement of ZAKα from the ribosome. Right: Model for sensing of a common ribotoxic stress signal by ZAKα based on mRNA stasis elicited by paused, stalled and collided ribosomes. In this model, activation of ZAKα is determined by prolonged and/or stable ribosome interaction. **f.** Cells from (a) were treated with torin (1 μM – 6 h) or increasing concentrations of the EEF2K-activating compound nelfinavir (10, 20, 40, 50 μM – 6 h). Lysates were analyzed as in (a).

### Ribosome-templated unfolding and transient dimerization underlies ZAKα kinase activation

Rather than a differential ribosome binding mode, sensing of mRNA stasis by ZAKα appears to be associated with a prolonged or more stable interaction (Fig. 4e) that sequesters the kinase in an activation-prone state. This model begs the question of how the ribosome increases the activation potential of a ZAKα molecule. ZAKα activation critically depends on activation loop auto-phosphorylation (*10, 39*), a process that most often occurs *in trans* (*40*). We thus tested whether forced dimerization of two ZAKα molecules led to kinase activation. To this end, we fused either full-length (FL) HA-tagged ZAKα or a construct containing only the kinase domain and the LZ (amino acids 1-332) with FKBP12-F36V. To enforce dimerization, we treated transfected cells with the bivalent FKBP12-binding compound AP20718 (Fig. 5a). For either construct, this led to an auto-phosphorylation-associated mobility shift of ZAKα (Fig. 5b,c). For the shorter construct, treatment with AP20718 even led to activation of both p38 and JNK (Fig. 5b), all in the absence of a ribotoxic stress signal. For the full-length construct, p38 and JNK activation was not observable (Fig. S4e), suggesting that the additional structured domains (compared to the 1-332 fragment) strongly impairs downstream signaling. While ZAKα appears to be a predominantly monomeric protein, it does have a propensity for self-interaction. We could thus co-purify transiently transfected GFP-tagged ZAKα and strep-HA-tagged ZAKα (Fig. S5a).

**Figure 5.**
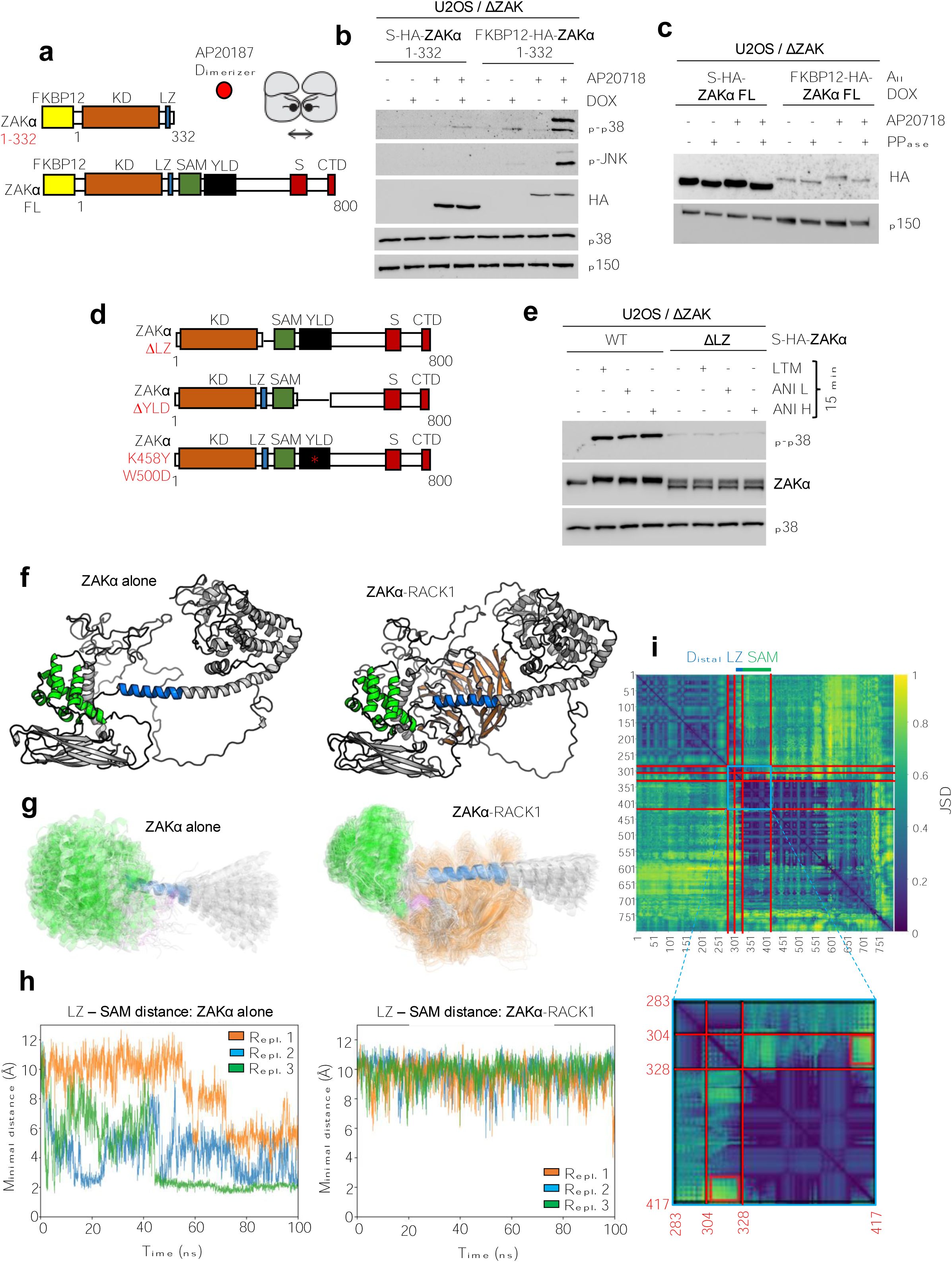
Ribosome-bound ZAKα is competent for dimerization and activation *in trans*. **a.** Experimental strategy for forced dimerization of ZAKα monomers. Full-length (FL) and truncated (aa 1-332) ZAKα were N-terminally fused to FKBP12-F36V, which can be forcibly dimerized using the bivalent small molecule binder AP20187. **b.** U2OS / ΔZAK cells conditionally expressing FKBP12-HA-ZAKα(1-332) from (a) were treated with doxycycline (DOX - overnight) and AP20187 (50 nM - 1 h) as indicated. Lysates were analyzed by immunoblotting with the indicated antibodies. **c.** U2OS / ΔZAK cells expressing full-length FKBP12-HA-ZAKα from (a) were treated with AP20187 (15 μM - 1 h) as indicated. Lysates were treated with 11 phosphatase (PPase) for 30 min at room temperature as indicated and analyzed as in (b). **d.** Schematic of ZAKα mutants with deletion of LZ (ΔLZ) and YLD (ΔYLD) and a composite point mutant of the YLD domain (K458Y, W500D). **e.** U2OS / ΔZAK cells stably rescued with WT and ΔLZ forms of strep-HA-tagged ZAKα were treated with ribotoxic stress agents anisomycin (ani L – 0.19 μM; ani H - 76 μM) or lactimidomycin (LTM – 1 μM) for 15 min and analyzed as in (b). **f.** Structure at t = 0 for ZAKα alone (left) and ZAKα-RACK1 (right) molecular dynamics (MD) simulations. ZAKα residues 336 to 417 (SAM domain - green), 309 to 329 (C-terminal part of LZ - blue) and RACK1 (orange) are highlighted. **g.** Structural ensemble samples from the ZAKα alone (left) and ZAKα-RACK1 (right) MD simulations, respectively, color-coded as in (f). In addition, ZAKα residues 418 to 422 (RACK1-binding SLIM - pink) are highlighted. For clarity, only residues 286 – 425 are shown, full-length ensembles are shown in Fig. S5k. Each ensemble shows 150 frames sampled at regular intervals from the concatenated triplicate simulations. **h.** Minimum distance in Angstroms (Å) between ZAKα residues 309 to 329 and residues 336 to 417 per replicate in the ZAKα alone (left) and ZAKα-RACK1 (right) MD simulations. **i.** Jensen-Shannon Divergence (JSD) between ZAKα residue-residue distances in ZAKα alone vs. ZAKα-RACK1 100 nanoseconds (ns) MD simulations. Each of the two conditions were simulated in triplicates, which were concatenated before calculating the JSD. Red lines indicate, from left to right (and top to bottom), residues 283, 304, 328 and 417. Red squares highlight JSD between ZAKα residues 400 to 417 and residues 304 to 328. Repl., replicates.

When presented with two ZAKα molecules, AF3 suggested a dimer conformation with the two kinase domains positioned face-to-face (consistent with activation loop *trans*-autophosphorylation) and guided by the two leucine zippers (Fig. S5b,c). It furthermore indicated that the YLD is a self-interacting domain that may stabilize or favor a dimerized state (Fig. S5b-d). We thus proceeded to delete the YLD in its entirety (ZAKα ΔYLD), yielding a mutant that was severely compromised for activation at 15 minutes after LTM, Ani Low and Ani High (Fig. 5d; Fig. S5e). At a later timepoint (1 hour), this mutant was however quite proficient in RSR signaling (Fig. S5f), suggesting a hypomorphic rather than null phenotype. Importantly, this deficiency was not associated with decreased ribosome binding as indicated by both sucrose cushions and iCLIP analysis (Fig. S5g,h). A similar conclusion was reached with a composite point mutant (ZAKα K458Y W500D) predicted (*41*) to abolish YLD domain dimerization (Fig. 5d; Fig. S5i). We also deleted the LZ in its entirety (ZAKα ΔLZ), and this mutant behaved as a null mutant despite being proficient in ribosome binding (Fig. 5d,e; Fig. S5j). We therefore conclude that the YLD is important, and that the LZ is critical, for translating ribotoxic stress sensing by ZAKα into kinase domain activation. These observations suggested that the two domains are required for establishing an activation-competent form of a dimeric or oligomeric complex. We previously reported that a patient-derived (ZAKα F368C (*42*)) as well as an engineered (ZAKα W347S) point mutation in the SAM domain were associated with constitutive ZAKα activity in the absence of an exogenous ribotoxic stress insult (*26*). In aggregate, all of these results imply that the YLD and LZ domains are positively-acting mediators of ZAKα dimerization and activation, while the SAM domain exerts negative regulation over this activity in the absence of ribosome binding and ribotoxic stress signals. One of the two SLIMs that ZAKα inserts into RACK1 upon ribosome binding is located right after the SAM domain, suggesting that this interaction (Fig. 1e-g) could restrain an inhibitory influence of the SAM domain on the neighboring and critical LZ. To test this, we conducted molecular dynamics simulations of full-length ZAKα either in isolation or complexed with RACK1 (in both cases, an AF3 model was used as the starting point (Fig. 5f)). We simulated in triplicates for 100 nanoseconds using 2 femtoseconds timesteps. Inspection of the resulting movies highlighted that the SAM vigorously explored the space around the distal part of the LZ, likely prohibiting the formation of a dimeric state that is competent for alignment of the two kinase domains (Movie S1). In the RACK1-complexed state, however, the SAM domain was effectively sequestered on the surface of RACK1 and restrained from influencing the LZ (Movie S2). The same conclusion was reached by inspecting structural ensembles of these movies. For simplicity of visual inspection, we overlaid only 150 of the frames and highlighted the ensembles by color-coding the locations of LZ and SAM (Fig. 5g; Fig. S5k). We furthermore computed the minimum distance between any residue in SAM and any residue in the distal (SAM-neighboring) part of the LZ over time from all simulation replicates. This analysis demonstrated the stochastic and transient nature of LZ-SAM proximity when ZAKα was simulated alone, a behavior that was completely obfuscated by binding to RACK1. (Fig. 5h). Finally, we computed the Jensen-Shannon Divergence (JSD) (*43*) between ZAKα residue-residue distances in ZAKα alone vs ZAKα-RACK1 simulations (Fig. 5i). This metric indicates the similarity between two probability distributions and also highlighted how the presence of RACK1 negatively affected the proximity over time of the distal part of LZ and SAM. In aggregate, our previous work, biochemical analyses and molecular dynamics simulations are consistent with a model in which intramolecular crosstalk between the negatively acting SAM domain and the essential LZ prohibits unscheduled ZAKα activation. It also suggests an explanation for how this auto-regulatory principle is bypassed by prolonged or stable binding of ZAKα to the ribosome.

## Discussion

Despite the overwhelming success of cryo-EM and cryo electron tomography (cryo-ET), these techniques have limitations when it comes to capturing very transient and/or low-stoichiometry complexes. In such cases, orthogonal approaches are warranted. Here, we used AF3 modelling (protein) combined with iCLIP (RNA) to unravel the ribosome-binding mode of the ribotoxic stress sensor ZAKα (Fig. 4e). Crucial to our result was the detection of a single dominant rRNA crosslink peak corresponding to the 18S helix-26. Of note, this result contrasts with that of our previous CLIP efforts, where we only found a weakly enriched peak corresponding to 18S helix-14 (*26*). In those studies, we transiently over-expressed a fragment encoding the C-terminal 100 amino acids of ZAKα. The present experiments are based on stable expression of full-length ZAKα which we now understand to engage in multiple, composite interactions with both rRNA and protein components of the ribosome. Regarding the two ribosomal binding patches for ZAKα, there are a couple of interesting parallels to known ribosome interactors. First, the critical binding to RACK1 appears to depend on a virtually identical interaction mode to that recently described for the mRNA stabilizing factors LARP4 and LARP4B (*31*). Thus, the SLIMs are highly similar between all three proteins and they all appear to occupy the same interaction clefts in RACK1. It is thus attractive to speculate that a number of proteins may contact RACK1 in a similar mode, giving rise to competition between factors and their associated translation surveillance activities. Second, the composite binding site comprised of RPS27 protein and the 18S helix-26 is also occupied by other factors than ZAKα. One of these is the large interaction factor EIF3 which both masks this site completely and blocks access to the mRNA exit channel (*44*). In addition, several IRES’es and the proteins USP10 and G3BP1 are known to bind the 18S helix-26 (*45, 46*). Of note, while RACK1 and both of its interactions with ZAKα are critical for RSR activation, this second binding site “only” confers optimal functionality in ribotoxic stress sensing. This is highlighted by the hypomorphic (but not non-functional) nature of a ZAKα mutant that cannot reach (ZAKα Δ625-760) from RACK1 to RPS27-18S helix-26.

Central to ribosome binding and ribotoxic stress sensing are the S and CTD domains of ZAKα. These are redundant mRNA binding entities as highlighted by the lack of crosslinking to mRNA (but still crosslinking to 18S helix-26 rRNA) by a composite mutant (ZAKα R->A ΔCTD). The CTD contains a large number of lysine and arginine residues that when mutated to alanines render this domain unfunctional. S domain function depends on the integrity of at least one of three peptide repeats with the consensus sequence **R**G**R**YXX**R/K** (*34*). These repeats are dispersed across the linker connecting the RACK1 and RPS27-helix-26 binding sites in the ZAKα C-terminus. We infer from the above that the positively charged amino acids in CTD as well as S (highlighted in bold - mutated to alanines in our mutant) connect to mRNA protruding from the ribosomal mRNA exit channel. In our bioinformatic analyses of iCLIP data, we did not find any sequence bias or enrichment of particular mRNA motifs. This is consistent with S and CTD being sequence-independent mRNA binders, similarly to what has been described for other intrinsically disordered regions (IDR) containing high contingents of positively charged amino acids (*47*).

To our knowledge there are no other examples of a similar translation stress-sensing mechanism. One close example is that of the *S. cerevisiae* E3 ubiquitin ligase Fap1 that contacts mRNA both at the entry and exit channels and therefore discriminates single stalled ribosomes from collision structures (*48*). Another example is that of the bacterial (*E. coli*) helicase HrpA that simultaneously loads onto static mRNA in the entry channel of a stalled ribosome and interacts with the collided ribosome entity of a disome (*23*). In this way, HrpA is a specific sensor of ribosome collision that uses ATP hydrolysis and helicase activity to split the structure. In both cases, Fap1 and HrpA act to dismantle stalled and collided ribosomes, respectively, while ZAKα acts upstream of a general stress signaling pathway.

It has been experimentally shown that ribosome collision results in stronger ZAKα activation than ribosome stalling (*16, 26, 49*). In light of the above, several explanations for this conundrum are possible. One possibility is that collisions simply provide for a more stably arrested template. Even high affinity inhibitors such as anisomycin, would dynamically vacate and re-bind the ribosomal A-site. Another possibility is that ZAKα is more prone to dimerize at a collision interface, given the close vicinity of additional low-affinity (mRNA-independent) binding sites on the stalled ribosome in a disome (Fig. S4d). Our work highlights that activation is associated with *trans*-autophosphorylation and thus obligatory self-interaction of ZAKα, most likely brought about by the stabilized binding of a ZAKα monomer to RACK1. Molecular dynamics simulations suggest that this interaction sequesters the SAM domain of ZAKα from exerting negative influence on the formation of an activation-competent dimer coordinated by the LZ domain. This activation mechanism resembles that of another mixed lineage kinase family member, MLK3. In the monomeric and auto-inhibited form of this kinase, the LZ is occluded by an intramolecular interaction between an SH3 domain and a proline residue (*50*). In the case of MLK3, the binding partner that disrupts this auto-inhibited state is GTP-bound Cdc42/Rac1 that can occupy a CRIB (Cdc42- and Rac-Interactive Binding) motif adjacent to the negatively acting proline. Once this intramolecular interaction is disrupted, the LZ of MLK3 mediates dimerization, trans-autophosphorylation and activation of MLK3 (*51*). Based on published data and the present work, we therefore propose that the SAM domain is the negatively acting feature in ZAKα and that RACK1 is the binding partner that can relieve this auto-inhibition.

In sum, our modelling and experimental validation places ZAKα at the mRNA exit channel where it monitors mRNA movement and thus ribosome processivity. By surveying this common feature of aberrant translation, ZAKα can detect pausing, stalling and collision through a common mechanism. We suggest that this insight unifies the disparate perceptions of stress sensing by the RSR-activating kinase ZAKα in the field of translation regulation and surveillance.

## Acknowledgements

We thank Drs Eric Bennett and Xiaoyun Lu for the kind gifts of reagents. We furthermore acknowledge the SUND Genomics Platform (University of Copenhagen, Denmark) for sequencing of iCLIP libraries. Work in the Bekker-Jensen lab was supported by the European Research Council (ERC) under the European Union’s Horizon 2020 research and innovation program (grant agreement 863911 - PHYRIST), Independent Research Fund Denmark (grant no. 3101-00344B) and the LEO Foundation (grant no. LF-OC-23-001458). Center for Gene Expression (CGEN) is a Center of Excellence funded by The National Danish Research Foundation (grant no. DNRF166). Work in the Heick Jensen lab was supported by the Novo Nordisk Foundation (ExoAdapt Grant 31199). A.B. was supported by the Marie Curie Individual Fellowship (EXOonRNA, 101026781) and Lundbeck Foundation Experiment Grant (R346-2020-1610).

## Declaration of interests

The authors have no positions, patents or financial interests to declare.

## Supplementary Movies

### Movie S1

Molecular dynamics simulation movie of ZAKα alone corresponding to Fig. 5f-h and Fig. S5k. ZAKα residues 336 to 417 (SAM domain - green) and 309 to 329 (C-terminal part of LZ - blue) are highlighted. Time frame: 100 ns.

### Movie S2

Molecular dynamics simulation movie of ZAKα-RACK1 corresponding to Fig. 5f-h and Fig. S5k. ZAKα residues 336 to 417 (SAM domain - green) and 309 to 329 (C-terminal part of LZ - blue) and RACK1 (orange) are highlighted. Time frame: 100 ns.

## Methods

### Plasmids

Plasmids containing truncations of ZAKα were PCR-cloned into pcDNA4/TO/Strep-HA using NotI restriction sites and small internal deletions were generated using an overlapping primer-based method (*52*). The W500D+K458Y mutant was synthetized by Integrated DNA Technologies (IDT) and subcloned into pcDNA4/TO/Strep-HA using NotI restriction sites. For generation of pcDNA4/TO/FKBP12-HA-ZAKα (FL and 1-332), FKBP12 was cloned into the vector at the expense of the strep-tag. gRNAs for CRIPSR/Cas9-mediated generation of HeLa ΔRACK1 cells were cloned using the pX459 plasmid (Addgene #62988). In brief, gRNA DNA oligos were ordered as complimentary sequences with overhangs and mixed at a 1:1 ratio for annealing. pX459 was digested with BbsI, and the gRNA was introduced using standard ligation reaction (New England Biolabs). The following gRNA sequences were used (*53*): RACK1-1-Fw; 5’-CACCGATTCCACAGCGTGCTCTTGCG, RACK1-1-Rv; 5’- AAACCGCAAGAGCACGCTGTGGAATC. For construction of the pX330A_sgC-EIF3A plasmid for endogenous degron tagging, gRNA oligos against the sequence 5’-atgagactgatgaagatgga (targeting the C-terminus of EIF3A) were annealed and cloned into BbsI-digested Cas9-encoding vector GW223-pX330A-sgX-sgPITCh, resulting in plasmid pX330A_sgC-EIF3A. For construction of pCRIS_PITChv2_Puro_Cterm_EIF3A, the FKBP12F36V-PuroR donor construct was generated by amplifying the FKBP12F36V sequence from pCRISPITChv2_Puro_BRD4_Nterm using primers containing EIF3A-specific microhomology sequences. The purified fragment was then Gibson assembled into the MluI-digested pCRIS_PITChv2_Puro_BRD4_Cterm backbone, yielding pCRIS_PITChv2_Puro_Cterm_EIF3A. All constructs were verified by sequencing, and all plasmid transfections were done using FUGENE6 (Promega) according to the manufacturer’s protocol.

### Cell culture and reagents

Female human osteosarcoma cells (U2OS) cells (ATCC, HTB-96; RRID: CVCL0042), female human malignant cervical epithelial cells (HeLa) (ATCC, CCL-2; RRID: CVCL0030) and female human embryonic kidney (HEK293) (ATCC, CRL-1573; RRID: CVCL0045) /EIF3A-HA-FKBP12 were cultured in Dulbecco’s Modified Eagle’s Medium (DMEM, Biowest # L0104-500) supplemented with a 10% fetal bovine serum, penicillin and streptomycin. Male human near haploid cells (HAP1) (RRID: CVCLY019) were cultured in Iscove’s Modified Dulbecco’s Medium (IMDM) GlutaMAX™ Supplement (Gibco # 31980030) supplemented with 10% FBS, penicillin, and streptomycin. All cells were cultured at 37°C in a humidified 5-8% CO2 cell incubator.

Derivative cell lines U2OS/ΔZAK, U2OS/ΔZAK/strep-HA-ZAKα_WT, U2OS/ΔZAK/strep-HA-ZAKα_R/K→A, U2OS/ΔZAK/strep-HA-ZAKα_ΔCTD, U2OS/ΔZAK/strep-HA-ZAKα_Δ670-712, U2OSΔZAK_R/K→A ΔCTD have been previously described (*26, 34*). HAP1/ΔRACK1 and HAP1/ΔRACK1/FLAG_RACK1 cells were a gift from Eric Bennett. To generate cell lines stably expressing truncations, internal deletions and point mutants of ZAKα under doxycycline inducible promoters, cells were co-transfected with pcDNA4/TO/Strep-HA-ZAKα constructs and pcDNA6/TR (Life Technologies) in a 1:4 ratio and selected for 14 days with zeocin (200 μg/ml) and blasticidin (5 μg/ml). Individual clones were picked, and expression analysed by immunofluorescence and western blotting. Unless otherwise indicated in figure legends, all U2OS/ΔZAK rescue cell lines were treated with doxycycline overnight to induce expression of the transgene, To generate HEK293/EIF3A-HA-FKBP12, the plasmids were co-transfected into HEK293 cells, and puromycin selection was applied to enrich for edited clones. Successful integration of the degron tag was validated by western blotting. Chemicals and inhibitors used in this study were: Doxycycline (Merck, D3347, 0.13 μg/ml, overnight), anisomycin (Merck, A9789), puromycin (Cayman Chemical, 13884), lactimidomycin (Merck, 5062910001), dTAG^V^-1 (Tocris, #6914), torin (InvivoGen, #inh-tor1), nelfinavir (MedChemExpress #HY-15287), AP20718 (MedChemExpress #HY-13992), ZAK inhibitor 6p (a gift from Xiaoyun Lu), zeocin (Thermo Fisher #R25001) and blastidicin (Thermo Fisher # R21001).

### Phosphatase treatment

Cells were lysed in EBC buffer without EDTA and phosphatase inhibitors and MnCl_2_ was added to a final concentration of 1 mM. 400 U of lambda phosphatase (New England Biolabs) were added or not, and the extracts were incubated for 30 min at 30 °C, directly mixed with Laemmli sample buffer, and boiled for 5 min before western blotting.

### Western blotting, pull-down and antibodies

For whole-cell extracts, cells were lysed in EBC buffer (50 mM Tris, pH 7.5, 150 mM NaCl, 1 mM EDTA, 0.5% NP-40, protease and phosphatase inhibitors), mixed with Laemmli sample buffer and boiled for 5-10 min. Pull-downs were done with Strep-Tactin Sepharose (IBA Life Sciences). Protein samples were resolved by SDS-PAGE and transferred to nitrocellulose membranes. Membranes were blocked in PBS-T + 5% milk before incubation with primary antibody overnight at 4 °C. Membranes were then washed in PBS-T and incubated with secondary antibody for 1 h at room temperature, before being washed in PBS-T and visualized by chemiluminescence (Clarity Western ECL substrate, Bio-Rad) using the Bio-Rad Chemidoc imaging system. Antibodies used: Rabbit polyclonal anti-ZAKα (Bethyl, Cat#A301-993A; RRID: AB_1576612), Mouse monoclonal anti-phospho-p38 (Cell Signaling, Cat#9216; RRID: AB_331296) Rabbit monoclonal anti-phospho-p38 (Cell Signaling, Cat#4511S; RRID: AB_2139682), Rabbit polyclonal antibody anti-p38 (Cell Signaling, Cat#9212; RRID: AB_330713), Mouse monoclonal anti-phospho-SAPK/JNK (Cell Signaling, Cat#9255; RRID: AB_2307321), Rabbit monoclonal anti-phospho-SAPK/JNK (Cell Signaling, Cat#4668, RRID:AB_823588), Rabbit monoclonal anti-SAPK/JNK (Cell Signaling, Cat#9258; RRID: AB_2141027), Rabbit polyclonal anti-ZAK (Proteintech, Cat#14945-1-AP; RRID: AB_1064269), Mouse monoclonal anti-p150 (BD biosciences, Cat#610473, RRID: AB_397845), Mouse monoclonal anti-α-Tubulin (Merck, Cat#T9026, RRID: AB_477593), Mouse monoclonal anti-HA-tag (Santa Cruz Biotechnology, Cat#sc-7392 HRP, RRID: AB_2894930), Mouse monoclonal anti-Puromycin (Millipore, Cat#MABE343, RRID: AB_2566826), Rabbit polyclonal anti-RACK1 (Bethyl, Cat#A302-545A, RRID:AB_1999012), Rabbit monoclonal-anti RPS10 (Abcam # ab151550, RRID:AB_2714147), Mouse monoclonal anti-GFP (Roche Cat# 11814460001, RRID:AB_390913), Mouse monoclonal anti-gamma-tubulin (Sigma, #T5326, RRID:AB_532292), Mouse monoclonal anti-RPL19 (Novus Biologicals #H00006143-M01, RRID:AB_509253).

### Sucrose cushions

Crude cellular ribosome pellets were generated by lysing cells in lysis buffer (15 mM Tris, pH 7.5, 0.5% NP40, 6 mM MgCl_2_, 300 mM NaCl, RiboLock RNase Inhibitor #E00381) and clearing the lysate at 12,000 g, 4 °C, 10 min. The supernatant was carefully layered onto a sucrose cushion (30% sucrose in 20 mM Tris, pH 7.5, 2 mM MgCl_2_, 150 mM KCl) and ultra-centrifuged at 38,800 rpm for 16 h using a Sorvall wX+ Ultrafuge and a FIBERlite F50L-8x39 rotor. Supernatants were discarded and pellets were washed in PBS and re-suspended (100 mM KCl, 5 mM MgCl_2_, 20 mM HEPES, pH 7.6, 1 mM DTT and 10 mM NH_4_Cl). Pellets and whole cell extracts were boiled in Laemmli buffer and analyzed by Western Blot.

### Polysome profiling

Cells were exposed to 500 J/m^2^ UVB as indicated. Following treatment, cytosolic lysates were prepared using 20 mM Hepes pH 7.5, 100 mM NaCl, 5 mM MgCl_2_, 100 μg/ml digitonin, 100 μg/ml cycloheximide, 1X protease inhibitor cocktail (Sigma, #P2714) and 200 U RiboLock RNase Inhibitor (Thermo Fisher Scientific, #EO0382). Extracts were pushed 10 times through a 26G needle and incubated on ice for 5 min prior to centrifugation at 17,000 g for 5 min at 4 °C. After adding CaCl_2_ to a final concentration of 1 mM, lysates were optionally digested with 500 U micrococcal nuclease (MNase) (New England Biolabs, #M0247) for 30 min at 22 °C (*49*). Digestion was terminated by adding 2 mM EGTA. Equivalent amounts of lysate (150 mg of undigested RNA or 180 mg of MNase-digested RNA) were resolved on 15-50% sucrose gradients by centrifugation at 38,000 rpm in a Sorvall TH64.1 rotor for 2.5 h at 4 °C. The gradients were analyzed using a Biocomp density gradient fractionation system with continuous monitoring of the absorbance at 260 nm.

### Alphafold modelling

The code for running AlphaFold 3 (*27*) was obtained from the AlphaFold 3 GitHub repository (https://github.com/google-deepmind/alphafold3, v3.0.1), while model weights were procured from Google upon request. The model was run locally in a NVIDIA Quadro RTX 8000 GPU with unified memory and memory preallocation enabled. Runs consisted of ZAK:possible interactor dimers. Possible interactors were all canonical protein isoforms, as per Uniprot, in the Reactome pathway R-HSA-72766 (*28*).

### UV crosslinking, extraction, and immunoprecipitation of crosslinked ZAKα-RNA complexes

Cell lysate preparation and immunoprecipitation (IP) were performed as described in (*54*) with minor modifications from (*55*). In brief, U2OS cells at 80% confluency were subjected to 254 nm (UVC) irradiation at a dose of 150 mJ/cm² using a STRATALINKER2000. Two 15 cm plates were used per IP, and experiments were run and sequenced in duplicate for each condition. Cells were harvested, lysed, and sonicated using Branson Digital Sonifier. Whole-cell extracts were treated with TURBO DNAse and RNAse I (1:500 dilution) prior to IP using MagStrep Tactin-XT beads (IBA, Cat. No. 2-5090-010). Protein-RNA complexes were subjected to high salt washes, including freshly added 2M Urea in the wash buffer, L3-App linkers were then ligated to 3’ ends of co-immunoprecipitated RNA, followed by ^32^P radiolabelling. To prevent UV-induced autophosphorylation and degradation of ZAKα, the ZAK inhibitor-6p was added at a final concentration of 10 μM in the lysis buffer and 25 μM in the hot PNK reaction mixture during the radiolabelling step. After separation by PAGE electrophoresis and wet transferring to a nitrocellulose membrane, radiolabeled ZAK-RNA complexes were then excised from the membranes and digested with proteinase K to remove ZAKα. RNA was subsequently precipitated by following the protocol from (*54*).

### iCLIP library preparation and sequencing

Libraries were prepared as described in (*54*) using barcoded adapters and pooled at equimolar concentrations prior to sequencing on an Illumina NextSeq 2000 platform. Eight libraries were pooled and applied to a P2-100 XLEAP chip+kit, returning app. 500 Mreads. Demultiplexing, adapter trimming, quality filtering and PCR-duplicate removal was performed as previously described (*56*).

### Library demultiplexing, mapping and quality control

Reads were aligned twice depending on their final usage using STAR v2.7.11 (*57*) as outlined in (*56*) with minor modifications. For rRNA crosslink mapping over a structure and rRNA peak visualization, multimapping reads were allowed and counted in all aligned locations, alignment was made to the RefSeq (*58*) GRCh38.p14 reference genome, and rRNA genes were found according to the corresponding RefSeq GTF annotation file. For all other applications (metagene profiling, biotype/UTR counting), multimapping reads were allowed but only counted in the top-scoring location, using the Ensembl (*59*) GRCh38.p14 reference genome. The Ensembl reference annotation was supplemented with the tRNAs detected from tRNAscan-SE for the human genome (*60*).

### Analysis of iCLIP data

Crosslink sites were identified and reads counted as described in (*56*). FeatureCounts(*61*) was used to count number of crosslinked features, while the metagene profiles were prepared with RiboMiner (*62*) and Plastid (*63*), considering, for each gene, only the longest transcript or the longest unambiguous stretch respectively, excluding transcripts with less than 50 reads mapped to them and normalizing to total number of crosslinks in the quantified region when necessary.

### Visualization of Alphafold scores and iCLIP values in ChimeraX

Alphafold 3 contact probability scores were calculated, for each possible interactor reference sequence, from the output contact probability matrices C_n_ for the 5 predicted models 0 to 4. First, a consensus contact probability matrix (CP_i,j_) was obtained by

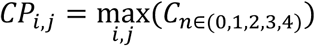

Then, an array representing the maximum contact probability (iCP) with chain A (ZAKα) for each residue in chain B (possible interactor) was defined as

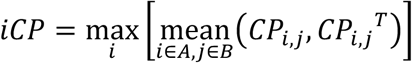

A corresponding reference sequence and array for the number of crosslinks per nucleotide was attained from the iCLIP data for the rRNA transcripts by directly mapping the number of crosslinks to each corresponding rRNA transcript at nucleotide resolution. These residue- and nucleotide-resolution values were then mapped from the reference protein and nucleotide sequences to those present in the target ribosome structure (4UG0) (*20*). Each sequence in the ribosome structure was assigned to the best aligning reference sequence. The iCP or number of crosslinks were transferred from reference to structure residues directly following the alignment, by mapping values from each reference residue to its aligned structure residue. The resulting arrays were then used to write a ChimeraX attribute file for visualization.

### Molecular dynamics simulation

Molecular dynamics simulations were run using GROMACS 2024.5 (*64*) in the CHARMM36 (*65*) force field. Simulation solvent consisted of 150 mM NaCl in water, plus any extra ions to reach net neutrality in the solution. The simulation pipeline consisted of an energy minimization step, followed by 5 ns NPT and 5 ns NVT equilibration, and finally a 100 ns production run, with 2 fs timesteps and a constant temperature of 310 K. RACK1 residues in direct contact with other ribosomal components (as per 4UG0) were restrained during production runs. We first ran the top ranked ZAKα-RACK1 Alphafold 3 structure once through the pipeline and used the output topology as input for all subsequent simulations. Both RACK1-present and -absent simulations were run in triplicates, generating random velocities for the atoms of each replicate once at the start of each NPT step. The initial topology for the RACK1-absent simulations was obtained by removing RACK1 from the input ZAKα-RACK1 and re-solvating ZAKα before starting the pipeline. Finally, simulation data was analysed using the Numpy (*66*), PENSA (*67*) (JSD calculation), and MDAnalysis (*68*) (inter-domain distance calculation) Python packages, while movies and superimpositions were composed in Pymol v3.1.

### Data and code availability

Demultiplexed fastq files containing iCLIP sequencing data, along with processed crosslinking BED files have been deposited in the NCBI Gene Expression Omnibus (GEO) with the accession number **GSE292064** and reviewer token **klmdcoiqdfknfqd**. The code used to map AF3 contact probability scores and iCLIP crosslinks to a structure is available at https://github.com/JoseFcoMH/defattr_painters.

## Notes

### Competing Interest Statement

The authors have declared no competing interest.

## References

1. A. C. Vind, F. L. Zhong, S. Bekker-Jensen, Death by ribosome. Trends Cell Biol, (2024).

2. M. S. Iordanov et al., Ultraviolet radiation triggers the ribotoxic stress response in mammalian cells. J Biol Chem 273, 15794–15803 (1998).

3. G. Snieckute et al., ROS-induced ribosome impairment underlies ZAKalpha-mediated metabolic decline in obesity and aging. Science 382, eadf3208 (2023).

4. M. S. Iordanov et al., Ribotoxic stress response: activation of the stress-activated protein kinase JNK1 by inhibitors of the peptidyl transferase reaction and by sequence-specific RNA damage to the alpha-sarcin/ricin loop in the 28S rRNA. Mol Cell Biol 17, 3373–3381 (1997).

5. N. J. Boon et al., DNA damage induces p53-independent apoptosis through ribosome stalling. Science 384, 785–792 (2024).

6. J. Xi et al., Initiation of a ZAKalpha-dependent ribotoxic stress response by the innate immunity endoribonuclease RNase L. Cell Rep 43, 113998 (2024).

7. A. Karasik, H. A. Lorenzi, A. V. DePass, N. R. Guydosh, Endonucleolytic RNA cleavage drives changes in gene expression during the innate immune response. Cell Rep 43, 114287 (2024).

8. A. C. Vind et al., The ribotoxic stress response drives acute inflammation, cell death, and epidermal thickening in UV-irradiated skin in vivo. Mol Cell 84, 4774–4789 e4779 (2024).

9. K. S. Robinson et al., ZAKalpha-driven ribotoxic stress response activates the human NLRP1 inflammasome. Science 377, 328–335 (2022).

10. N. K. Sinha et al., The ribotoxic stress response drives UV-mediated cell death. Cell, (2024).

11. A. Subramanian et al., A Legionella toxin exhibits tRNA mimicry and glycosyl transferase activity to target the translation machinery and trigger a ribotoxic stress response. Nat Cell Biol 25, 1600–1615 (2023).

12. C. McKenney et al., CDK4/6 activity is required during G(2) arrest to prevent stress-induced endoreplication. Science 384, eadi2421 (2024).

13. A. C. Vind, G. Snieckute, S. Bekker-Jensen, M. Blasius, Run, Ribosome, Run: From Compromised Translation to Human Health. Antioxid Redox Signal 39, 336–350 (2023).

14. P. W. Ford, M. Narasimhan, E. J. Bennett, Ubiquitin-dependent translation control mechanisms: Degradation and beyond. Cell Rep 43, 115050 (2024).

15. J. Cordes, S. Zhao, C. M. Engel, J. Stingele, Cellular responses to RNA damage. Cell 188, 885–900 (2025).

16. C. C. Wu, A. Peterson, B. Zinshteyn, S. Regot, R. Green, Ribosome Collisions Trigger General Stress Responses to Regulate Cell Fate. Cell 182, 404–416 e414 (2020).

17. T. Inada, R. Beckmann, Mechanisms of Translation-coupled Quality Control. J Mol Biol 436, 168496 (2024).

18. M. B. D. Muller et al., The ribosome as a platform to coordinate mRNA decay. Nucleic Acids Res 53, (2025).

19. A. A. Pochopien et al., Structure of Gcn1 bound to stalled and colliding 80S ribosomes. Proc Natl Acad Sci U S A 118, (2021).

20. M. Narita et al., A distinct mammalian disome collision interface harbors K63-linked polyubiquitination of uS10 to trigger hRQT-mediated subunit dissociation. Nat Commun 13, 6411 (2022).

21. K. Q. Kim et al., Multiprotein bridging factor 1 is required for robust activation of the integrated stress response on collided ribosomes. Mol Cell 84, 4594–4611 e4599 (2024).

22. E. N. Park et al., B. subtilis MutS2 splits stalled ribosomes into subunits without mRNA cleavage. EMBO J 43, 484–506 (2024).

23. A. Campbell et al., The RNA helicase HrpA rescues collided ribosomes in E. coli. Mol Cell 85, 999–1007 e1007 (2025).

24. K. Saito et al., Ribosome collisions induce mRNA cleavage and ribosome rescue in bacteria. Nature 603, 503–508 (2022).

25. G. Snieckute et al., Ribosome stalling is a signal for metabolic regulation by the ribotoxic stress response. Cell Metab 34, 2036–2046 e2038 (2022).

26. A. C. Vind et al., ZAKalpha Recognizes Stalled Ribosomes through Partially Redundant Sensor Domains. Mol Cell 78, 700–713 e707 (2020).

27. J. Abramson et al., Accurate structure prediction of biomolecular interactions with AlphaFold 3. Nature 630, 493–500 (2024).

28. M. Milacic et al., The Reactome Pathway Knowledgebase 2024. Nucleic Acids Res 52, D672–D678 (2024).

29. H. Khatter, A. G. Myasnikov, S. K. Natchiar, B. P. Klaholz, Structure of the human 80S ribosome. Nature 520, 640–645 (2015).

30. M. Brockmann et al., Genetic wiring maps of single-cell protein states reveal an off-switch for GPCR signalling. Nature 546, 307–311 (2017).

31. A. Ranjan et al., The short conserved region-2 of LARP4 interacts with ribosome-associated RACK1 and promotes translation. Nucleic Acids Res 53, (2025).

32. J. S. Xiang, D. M. Schafer, K. L. Rothamel, G. W. Yeo, Decoding protein-RNA interactions using CLIP-based methodologies. Nat Rev Genet 25, 879–895 (2024).

33. J. Fedry et al., Visualization of translation reorganization upon persistent ribosome collision stress in mammalian cells. Mol Cell 84, 1078–1089 e1074 (2024).

34. V. B. I. Johansen, G. Snieckute, A. C. Vind, M. Blasius, S. Bekker-Jensen, Computational and Functional Analysis of Structural Features in the ZAKalpha Kinase. Cells 12, (2023).

35. N. Ahmed et al., Identifying A- and P-site locations on ribosome-protected mRNA fragments using Integer Programming. Sci Rep 9, 6256 (2019).

36. G. Leprivier et al., The eEF2 kinase confers resistance to nutrient deprivation by blocking translation elongation. Cell 153, 1064–1079 (2013).

37. A. De Gassart et al., Pharmacological eEF2K activation promotes cell death and inhibits cancer progression. EMBO Rep 17, 1471–1484 (2016).

38. P. R. Smith et al., eEF2K regulates pain through translational control of BDNF. Mol Cell 85, 756–769 e755 (2025).

39. E. Tosti, L. Waldbaum, G. Warshaw, E. A. Gross, R. Ruggieri, The stress kinase MRK contributes to regulation of DNA damage checkpoints through a p38gamma-independent pathway. J Biol Chem 279, 47652–47660 (2004).

40. R. Reinhardt, T. A. Leonard, A critical evaluation of protein kinase regulation by activation loop autophosphorylation. Elife 12, (2023).

41. Y. Zhou, Q. Pan, D. E. V. Pires, C. H. M. Rodrigues, D. B. Ascher, DDMut: predicting effects of mutations on protein stability using deep learning. Nucleic Acids Res 51, W122–W128 (2023).

42. M. Spielmann et al., Exome sequencing and CRISPR/Cas genome editing identify mutations of ZAK as a cause of limb defects in humans and mice. Genome Res 26, 183–191 (2016).

43. F. Nielsen, On a Generalization of the Jensen-Shannon Divergence and the Jensen-Shannon Centroid. Entropy (Basel) 22, (2020).

44. J. Brito Querido et al., The structure of a human translation initiation complex reveals two independent roles for the helicase eIF4A. Nat Struct Mol Biol 31, 455–464 (2024).

45. H. Yamamoto et al., Molecular architecture of the ribosome-bound Hepatitis C Virus internal ribosomal entry site RNA. EMBO J 34, 3042–3058 (2015).

46. C. Meyer, A. Garzia, P. Morozov, H. Molina, T. Tuschl, The G3BP1-Family-USP10 Deubiquitinase Complex Rescues Ubiquitinated 40S Subunits of Ribosomes Stalled in Translation from Lysosomal Degradation. Mol Cell 77, 1193–1205 e1195 (2020).

47. A. Zeke et al., Deep structural insights into RNA-binding disordered protein regions. Wiley Interdiscip Rev RNA 13, e1714 (2022).

48. S. Li et al., Sensing of individual stalled 80S ribosomes by Fap1 for nonfunctional rRNA turnover. Mol Cell 82, 3424–3437 e3428 (2022).

49. M. Stoneley et al., Unresolved stalled ribosome complexes restrict cell-cycle progression after genotoxic stress. Mol Cell 82, 1557–1572 e1557 (2022).

50. H. Zhang, K. A. Gallo, Autoinhibition of mixed lineage kinase 3 through its Src homology 3 domain. J Biol Chem 276, 45598–45603 (2001).

51. C. Rattanasinchai, K. A. Gallo, MLK3 Signaling in Cancer Invasion. Cancers (Basel) 8, (2016).

52. H. Liu, J. H. Naismith, An efficient one-step site-directed deletion, insertion, single and multiple-site plasmid mutagenesis protocol. BMC Biotechnol 8, 91 (2008).

53. S. Jha et al., Trans-kingdom mimicry underlies ribosome customization by a poxvirus kinase. Nature 546, 651–655 (2017).

54. A. Buchbender et al., Improved library preparation with the new iCLIP2 protocol. Methods 178, 33–48 (2020).

55. R. A. Cordiner et al., Temporal-iCLIP captures co-transcriptional RNA-protein interactions. Nat Commun 14, 696 (2023).

56. A. Busch, M. Bruggemann, S. Ebersberger, K. Zarnack, iCLIP data analysis: A complete pipeline from sequencing reads to RBP binding sites. Methods 178, 49–62 (2020).

57. A. Dobin et al., STAR: ultrafast universal RNA-seq aligner. Bioinformatics 29, 15–21 (2013).

58. N. A. O’Leary et al., Reference sequence (RefSeq) database at NCBI: current status, taxonomic expansion, and functional annotation. Nucleic Acids Res 44, D733–745 (2016).

59. S. C. Dyer et al., Ensembl 2025. Nucleic Acids Res 53, D948–D957 (2025).

60. P. P. Chan, T. M. Lowe, tRNAscan-SE: Searching for tRNA Genes in Genomic Sequences. Methods Mol Biol 1962, 1–14 (2019).

61. Y. Liao, G. K. Smyth, W. Shi, featureCounts: an efficient general purpose program for assigning sequence reads to genomic features. Bioinformatics 30, 923–930 (2014).

62. F. Li, X. Xing, Z. Xiao, G. Xu, X. Yang, RiboMiner: a toolset for mining multi-dimensional features of the translatome with ribosome profiling data. BMC Bioinformatics 21, 340 (2020).

63. J. G. Dunn, J. S. Weissman, Plastid: nucleotide-resolution analysis of next-generation sequencing and genomics data. BMC Genomics 17, 958 (2016).

64. S. Pronk et al., GROMACS 4.5: a high-throughput and highly parallel open source molecular simulation toolkit. Bioinformatics 29, 845–854 (2013).

65. J. Huang et al., CHARMM36m: an improved force field for folded and intrinsically disordered proteins. Nat Methods 14, 71–73 (2017).

66. C. R. Harris et al., Array programming with NumPy. Nature 585, 357–362 (2020).

67. M. Vogele et al., Systematic analysis of biomolecular conformational ensembles with PENSA. J Chem Phys 162, (2025).

68. N. Michaud-Agrawal, E. J. Denning, T. B. Woolf, O. Beckstein, MDAnalysis: a toolkit for the analysis of molecular dynamics simulations. J Comput Chem 32, 2319–2327 (2011).

