## Supplementary Figures and Legends for "ZAKα is a sensor of mRNA stasis at the ribosomal exit channel"

Vind et al, Figure S1

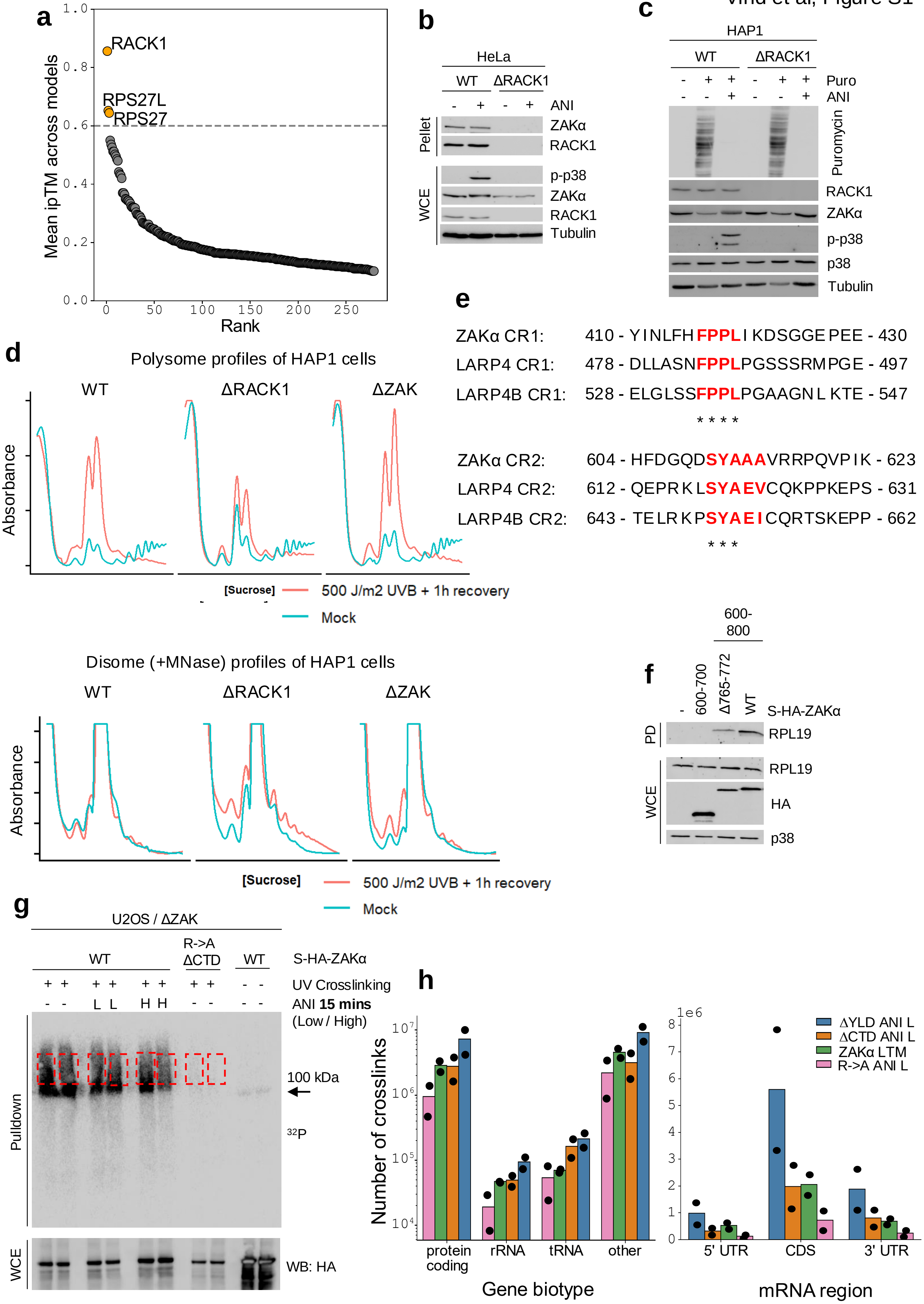

**Figure S1.**

**RACK1 is required for ZAKα-ribosome interaction**

**a.** Alphafold3 (AF3) prediction scores for ZAKα binding to proteins in the Reactome (reactome.org) pathway "Translation", R-HSA-72766. Predictions are ranked according to ipTM score and dashed line indicates our cutoff score of ipTM = 0.6. **b.** Hela WT and ΔRACK1 cells were treated with ani (1 μM - 1 h) and lysates were ultracentrifuged through a sucrose cushion. Whole cell extract (WCE) and pelleted material (pellet) containing ribosomes were analyzed by immunoblotting with the indicated antibodies. **c.** HAP1 WT and ΔRACK1 cells were treated with anisomycin (0.5 μg/ml – 1 hour). Puromycin (10 μg/ml) was added to the culture 10 min prior to harvest and lysates were analyzed by immunoblotting with the indicated antibodies. **d.** Cells from (b) were irradiated with UVB (500 J/$m^2$ - 1 hour). Lysates were treated (bottom) or not (top) with MNase to convert polysomes to ribosomes. Materials were separated on a linear sucrose gradient and UV absorbance was measured with a fraction collector to indicate RNA (ribosome) content. **e.** Alignment of ZAKα short linear interaction motifs (SLIMs) binding to RACK1 with motifs in LARP4 and LARP4B (CR1 and CR2) previously shown to occupy the same binding sites on RACK1. Note the high identity / similarity between residues that directly contact RACK1. **f.** U2OS cells were transfected with the indicated strep-HA tagged C-terminal ZAKα fragments. Lysates were subjected to strep purification and pull-down (PD) material and whole cell extract (WCE) were analyzed by immunoblotting with the indicated antibodies. **g.** U2OS / ΔZAK cells stably rescued with WT and ribosome-binding deficient (R->A ΔCTD) forms of strep-HA-tagged ZAKα were treated with ZAK inhibitor (10 μM – 30 min) and anisomycin (ani L – 0.19 μM; Ani H - 76 μM) for 15 min as indicated. Cells were crosslinked by UVC irradiation (150 mJ/$cm^2$) and lysates were treated with DNase and RNase and subjected to strep purification. RNA 3' ends were ligated to L3-App linkers and radiolabelled with $^{32}P$. Material was separated by PAGE and developed by autoradiography. Red boxes indicate the areas of the gel that were excised and processed for library preparation and sequencing. **h.** Left: Total number of ZAKα crosslinks from biological duplicate samples of sequencing batch 2 from Fig. 2a according to RNA category. Right: As in (left), except according to mRNA elements.

Vind et al, Figure S2

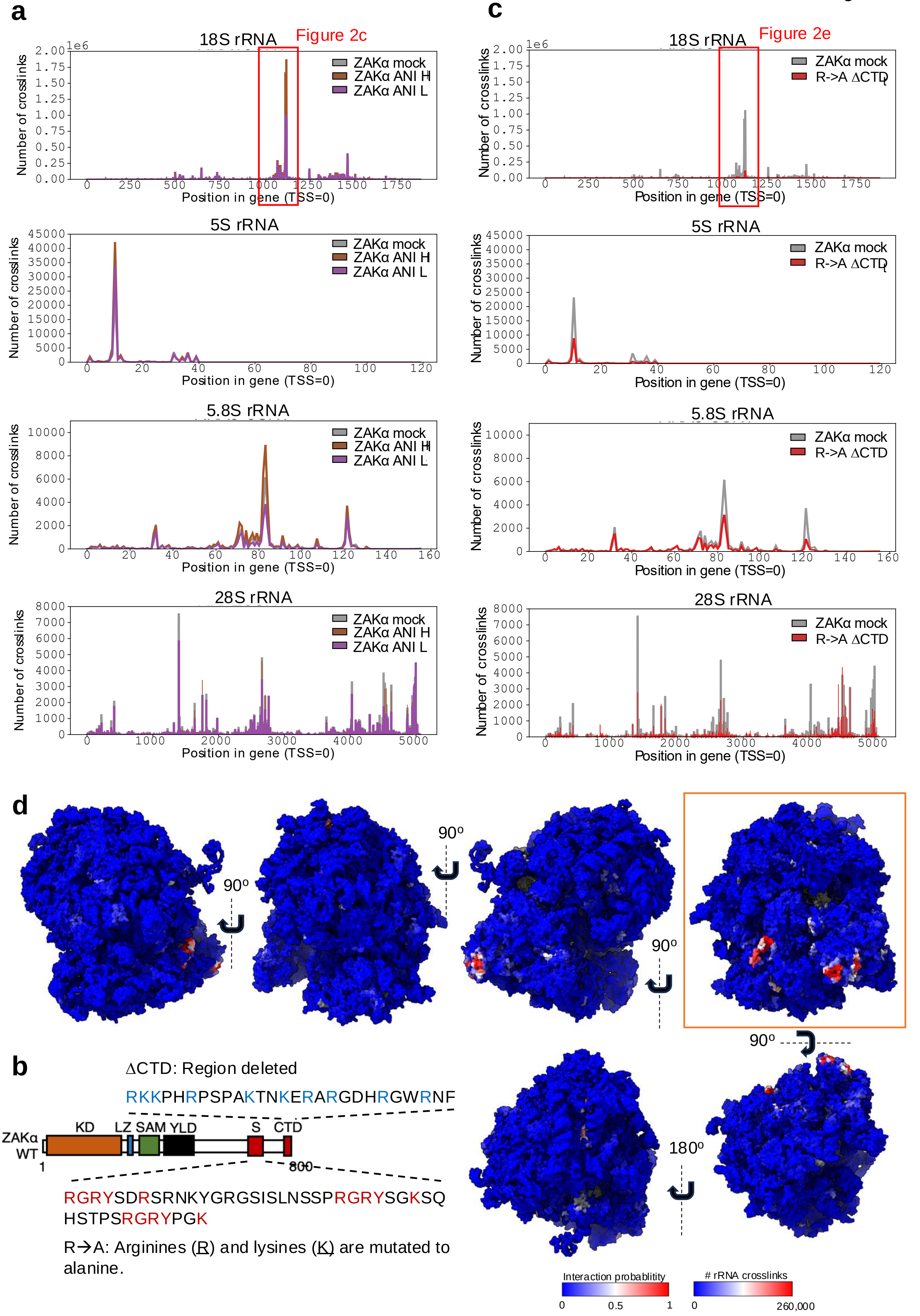

**Figure S2.**

**iCLIP highlights a single prominent rRNA interaction site for ZAKα**

**a.** Total number of sequenced ZAKα crosslinks for individual nucleotides across the four rRNAs. A single peak (red box) was orders of magnitude more abundant than any other peak. Mock treatment, low anisomycin (ani L) and high anisomycin (ani H) (15 min) conditions have been overlaid. Values represent the mean from duplicate experiments. **b.** Schematic of ZAKα domain composition. The CTD consists of a span of positively charged amino acids (R, K). The S domain contains three peptide repeats of similar sequence. Mutation of Rs and Ks (red) to As in the S domain combined with deletion of the CTD gives rise to the activation- and ribosome binding-deficient mutant of ZAKα (R->A ΔCTD). KD, kinase domain; LZ, leucine zipper; SAM, sterile alpha-motif; YLD, Yeats-like domain; S, sensor domain; CTD, C-terminal domain. **c.** As in (a), except that crosslinks from WT and R->A ΔCTD ZAKα have been overlaid. **d.** Structure of the human ribosome (PDB 4UG0) painted by per-residue ZAKα interaction probabilities and number of ZAKα crosslinks per rRNA residue. Pictures represent the structure from Fig. 2g viewed from all six sides, highlighting two likely ZAKα interaction sites on the ribosome.

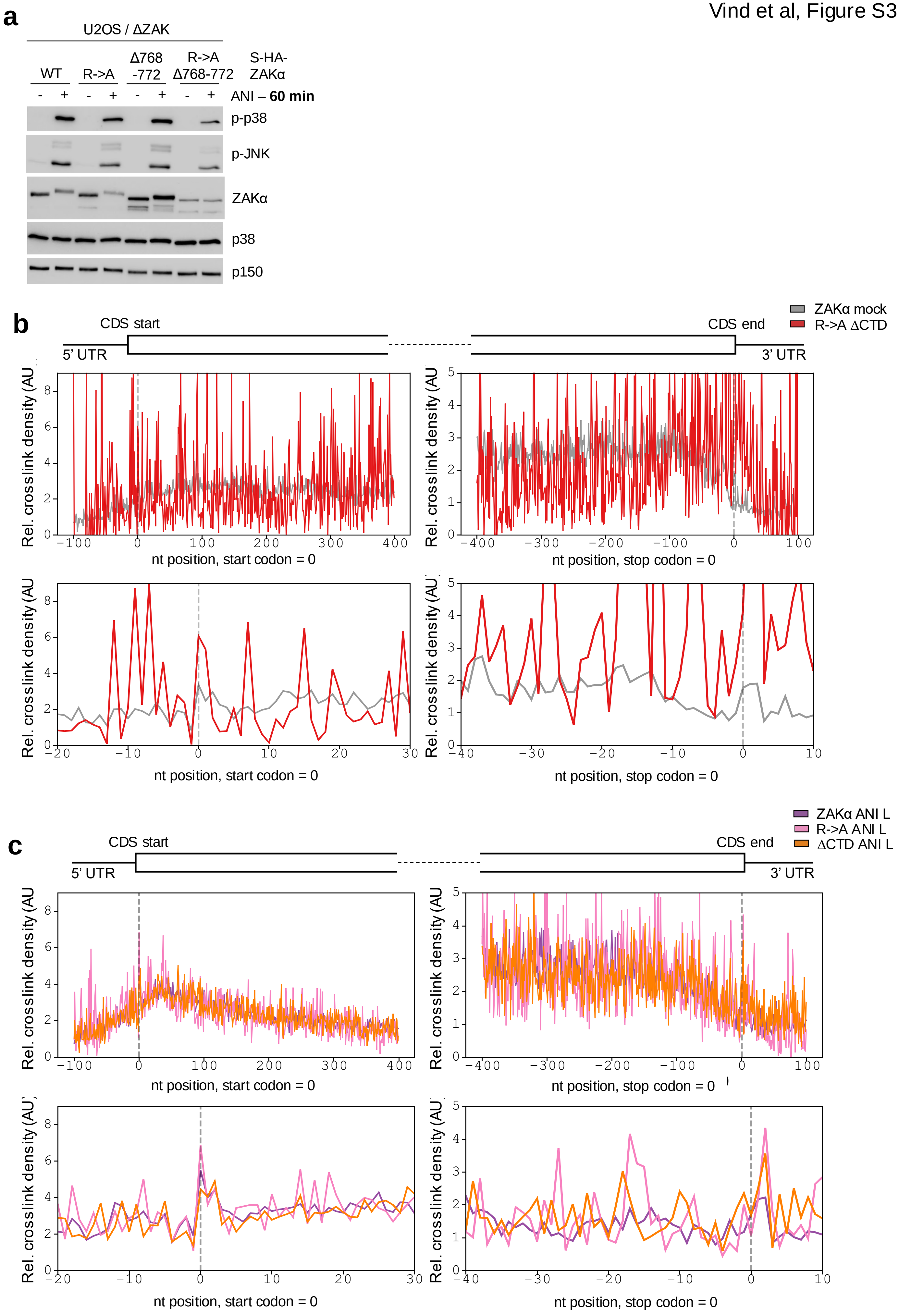
a
U2OS / ΔZAK
WT
R->A
Δ768 -772
R->A Δ768-772
S-HA-ZAKα
ANI – 60 min
p-p38
p-JNK
ZAKα
p38
p150
b
CDS start
5' UTR
CDS end
3' UTR
ZAKα mock
R->A ΔCTD
Rel. crosslink density (AU)
nt position, start codon = 0
nt position, stop codon = 0
c
ZAKα ANI L
R->A ANI L
ΔCTD ANI L

**Figure S3.**

**mRNA binding capability of ZAKα mutants**

**a.** U2OS / ΔZAK cells stably rescued with strep-HA-tagged ZAKα mutants deficient for RPS27 binding and/or S domain functionality were treated with anisomycin (ani, 1 μM – 60 min) and analyzed by immunoblotting with the indicated antibodies. **b.** Analysis of normalized mRNA crosslinks for WT and R->A ΔCTD ZAKα around the start (left) and stop codons (right) shown at low (top) and high (bottom) resolution. Values represent the mean from duplicate experiments. **c.** As in (b), except that cells expressing WT, R->A and ΔCTD forms of strep-HA-tagged ZAKα were treated with ani (1 μM – 15 min). nt, nucleotide number.

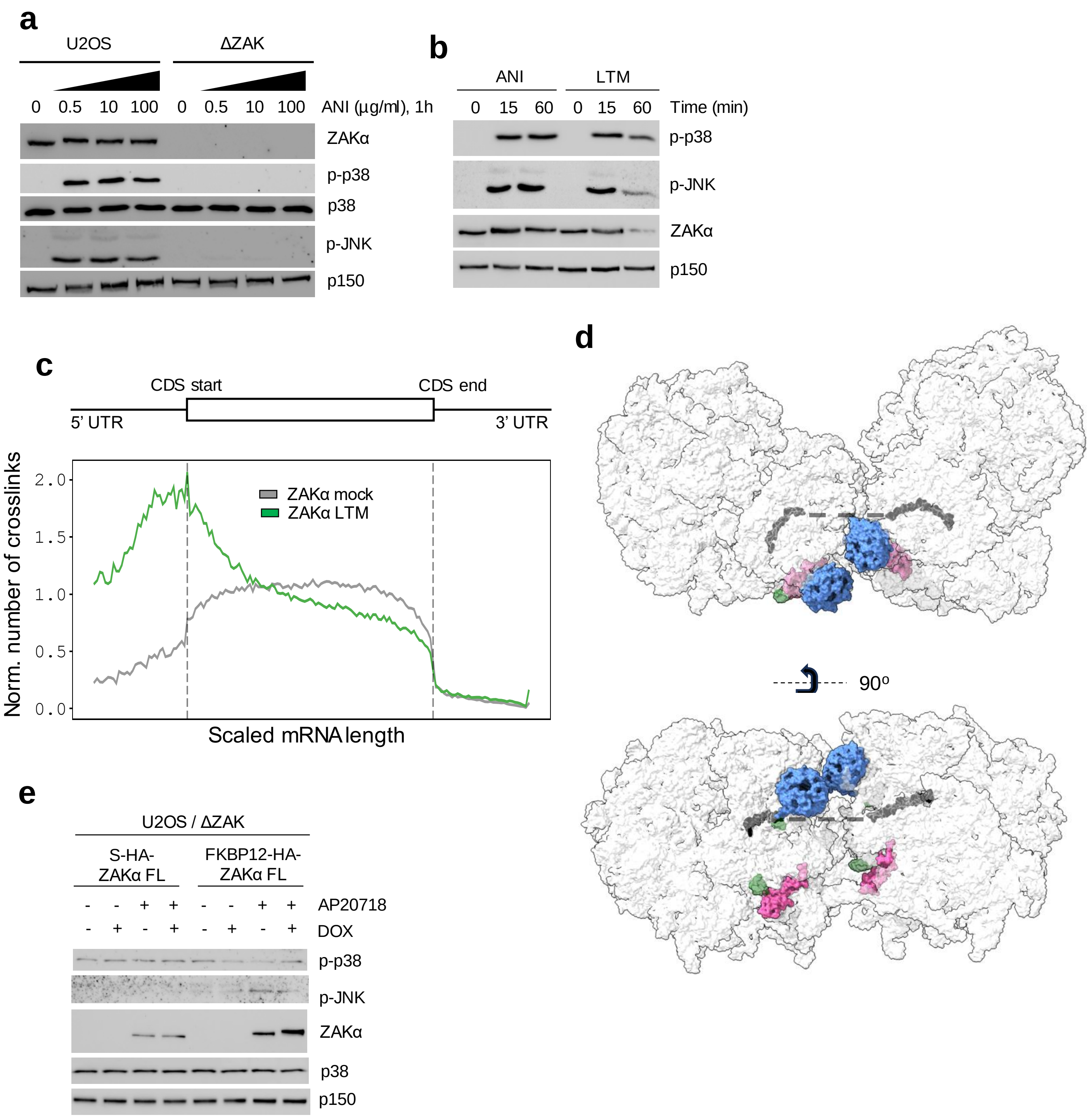
a
U2OS
ΔZAK
0 0.5 10 100 0 0.5 10 100 ANI (µg/ml), 1h
ZAKα
p-p38
p38
p-JNK
p150
b
ANI
LTM
0 15 60 0 15 60 Time (min)
p-p38
p-JNK
ZAKα
p150
c
CDS start
CDS end
5' UTR
3' UTR
ZAKα mock
ZAKα LTM
Norm. number of crosslinks
2.0
1.5
1.0
0.5
0.0
Scaled mRNA length
d
90º
e
U2OS / ΔZAK
S-HA-ZAKα FL
FKBP12-HA-ZAKα FL
- - + + - - + + AP20718
- + - + - + - + DOX
p-p38
p-JNK
ZAKα
p38
p150

**Figure S4.**

**ZAKα activation is independent of ribosome collision**

**a.** U2OS and ΔZAK cells were treated with increasing concentrations of anisomycin (ani – 1, 10 and 100 μg/ml) for 60 min. Lysates were analyzed by immunoblotting with the indicated antibodies. **b.** U2OS cells were treated with ribotoxic stress agents ani (1 μM) and lactimidomycin (LTM – 1 μM) for the indicated times. Lysates were analyzed as in (a). **c.** Metagene profiles of total number of crosslinks for mock and LTM-treated (1 μM, 15 min) WT ZAKα along scaled length of spliced mRNAs determined by iCLIP. Values represent the mean from duplicate experiments. **d.** Representation of a collided human ribosome structure (PDB 7QVP) with RACK1 (blue), RPS27 (magenta), 18S-helix26 (green) and mRNA (grey) highlighted. **e.** U2OS / ΔZAK cells conditionally expressing FKBP12-HA-ZAKα full-length (FL) from Fig. 5a were treated with doxycycline (DOX - overnight) and AP20187 (50 nM - 1 h) as indicated. Lysates were analyzed by immunoblotting with the indicated antibodies.

Vind et al, Figure S5

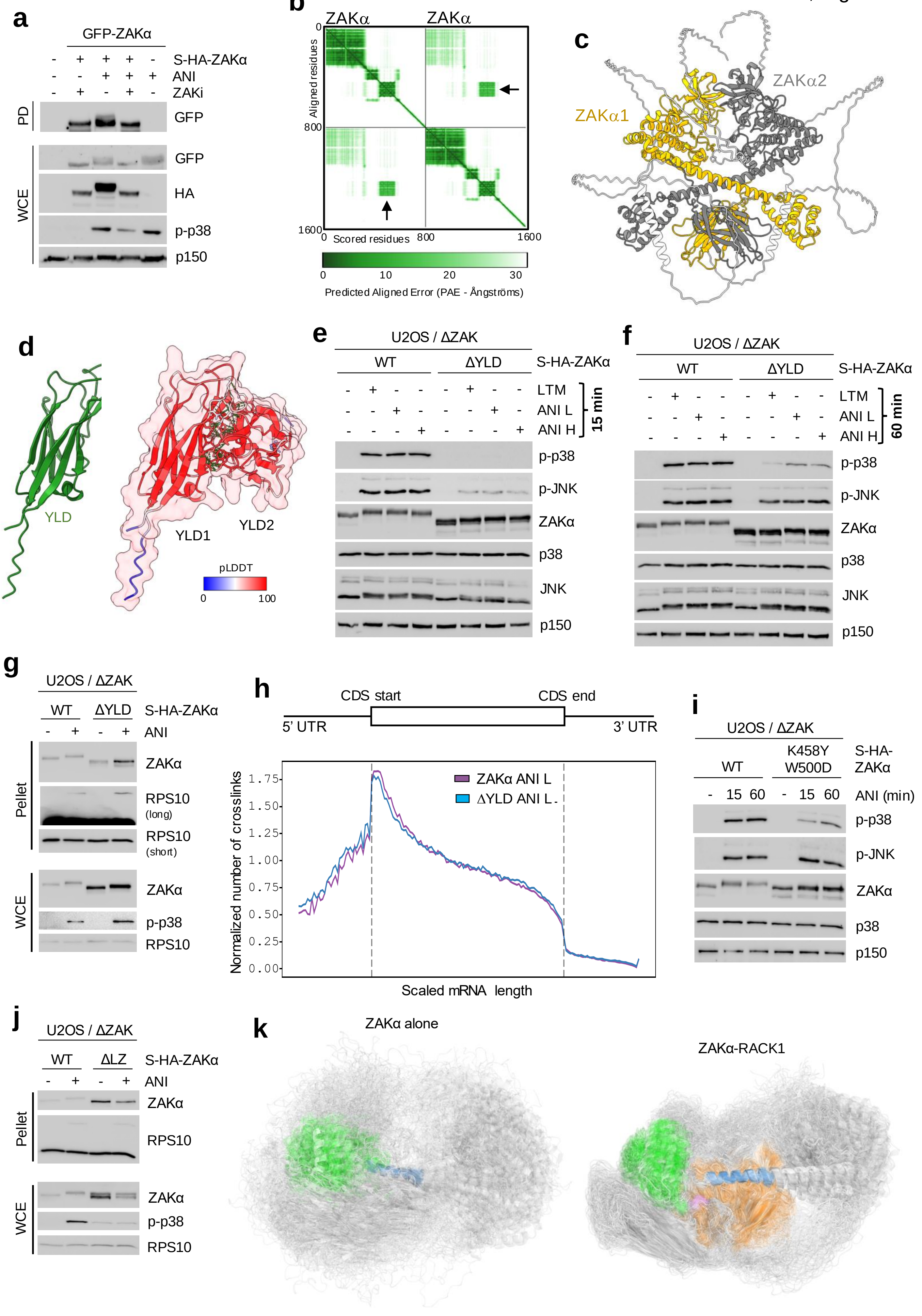

**Figure S5.**

**ZAKα YLD is required for optimal kinase activation**

**a.** U2OS cells stably expressing GFP-ZAKα were transfected with Strep-HA-ZAKα and treated with anisomycin (ani - 1 μM) and ZAK inhibitor (ZAKi - 1 μM) for 60 min as indicated. Lysates were subjected to strep purification and pull-down (PD) material and whole cell extract (WCE) were analyzed by immunoblotting with the indicated antibodies. **b.** Predicted aligned error (PAE) matrix plot of AF3-generated prediction of a ZAKα dimer. Values for YLD-YLD binding are indicated with black arrows. **c.** AF3-generated structure from (b). Sequences from start until end of the YLD domain are highlighted by color. **d.** Left: Predicted structure of an isolated ZAKα YLD domain. Right: Predicted structure of a ZAKα YLD dimer. Hydrogen bonds between chains are indicated and chains and interacting side chains are colored according to the predicted local distance difference test (pLDDT) score. **e.** U2OS / ΔZAK cells stably rescued with WT and ΔYLD forms of strep-HA-tagged ZAKα were treated with ribotoxic stress agents ani (L – 0.19 μM; H - 76 μM) or lactimidomycin (LTM – 1 μM) for 15 min and analyzed by immunoblotting with the indicated antibodies. **f.** As in (e), except that cells were treated with ani and LTM for 60 min. **g.** Cells from (e) were treated with ani (1 μM - 1 h) and lysates were ultracentrifuged through a sucrose cushion. Whole cell extract (WCE) and pelleted material (pellet) enriched for ribosomes were analyzed by immunoblotting with the indicated antibodies. **h.** Metagene profiles of total number of crosslinks for ani-treated (1 μM – 15 min) ZAKα WT and ΔYLD along scaled length of spliced mRNAs determined by iCLIP. Values represent the mean from duplicate experiments. **i.** U2OS / ΔZAK cells stably rescued with WT and YLD point mutated (K458Y W500D) forms of strep-HA-tagged ZAKα were treated with ani (1 μM) for the indicated times. Lysates were analyzed as in (e). **j.** As in (g), except that WT was compared to ΔLZ ZAKα. **k.** Full-length ensembles corresponding to (and colorcoded as in) Fig. 5g.
